# Plasma membrane order maps functional diversity in immune cells

**DOI:** 10.1101/2024.01.15.575649

**Authors:** Luca A. Andronico, Cenk O. Gurdap, Abishek Arora, Franziska Ragaller, Patrick A. Sandoz, Yidan Jiang, Sarantis Giatrellis, Leonard L. de Boer, Valentina Carannante, Sofiia Iskrak, Jaromir Mikes, Marcus Buggert, Anders Österborg, Björn Önfelt, Andrey Klymchenko, Petter Brodin, Erdinc Sezgin

## Abstract

Cell membranes undergo biophysical remodelling as an adaptation to the surroundings and to perform specific biological functions. However, the extent and relevance of such changes in human immune cells remain unknown, largely due to the lack of single-cell and multidimensional methodologies. Here, we apply a cytometry-based method to fill this gap by combining biophysical profiling with simultaneous analysis of immune cell markers. This platform reveals notable cell type-dependent plasma membrane order heterogeneity in immune cells. By sorting immune cells according to their membrane order and performing transcriptome and spatial surface proteome analyses together with functional tests, we show that plasma membrane order can be used to identify subsets of immune cells with distinct phenotypes and functional behaviours. Our findings demonstrate a broad heterogeneity of plasma membrane order in immune cells that will provide a more precise definition of immune cell states based on their biophysical properties in health and disease.

## Background

Immune cells are dynamically regulated across tissues, over the course of life and in relation to stimulation, offering spatiotemporal control of immunity. Recent data suggests a continuum of archetypal cell states as a key component of immune system heterogeneity^1^. This concept proposes that cells can assume various states depending on their microenvironment and stimulants therein^2^. While cell types are mostly defined by surface markers or transcriptional regulation and hence can be robustly distinguished^3^, cell state definitions are more complex, and requires multiparametric analysis of macromolecular content.

Lipids are fundamental to maintaining cell homeostasis as they allow for energy storage via lipid droplets formation^4^, organelle compartmentalization, by self-organizing in membranes^5,6^, and intracellular signalling by acting as secondary messengers^7^. Lipids exert their function by operating as a complex dynamic system of hundreds of molecules interacting with one another. This is evident in cell membranes, where the collective interaction between different lipid species dictates properties of the plasma membrane (PM) such as polarity, order (also referred to as membrane fluidity), viscosity, bending rigidity etc^8,9^. These biophysical properties have been shown to change during cellular processes such as proliferation and migration^10–14^. More recently, evidence has started highlighting the relationship between membrane biophysical properties and immune cell activity^15–23^. Importantly, alterations in cellular biophysical properties could occur without any changes in the genomic and transcriptomic profiles of dysfunctional cells (e.g., induced by nutrients or immediate microenvironment)^24,25^, which makes them complementary cell state indicators.

Cellular biophysical properties are commonly investigated by means of environment-sensitive fluorescent probes, whose photophysical properties change upon physical/chemical changes in the environment^26–31^. In particular, solvatochromic dyes, such as Laurdan, Nile Red and their derivatives, which shift their emission spectra as a function of local polarity and hydration, reflecting lipid order in biomembranes are widely used^32^. However, the use of such probes remains largely confined to microscopy techniques^33^, with only a few examples of their application in high-throughput methodologies such as flow cytometry. For instance, flow cytometry combined with environment-sensitive probes has been used to measure PM order in cell lines^34^ and bacteria^35–37^. Similar approaches have been applied to characterize PM order remodelling in immune cells. Miguel et al., for instance, focused on CD4⁺ T cells, comparing PM order differences between healthy donors and patients with autoimmune rheumatic disease^38^, demonstrating a strong correlation between membrane order and cell functionality. These studies focused on a specific cell type; however, capturing the broader heterogeneity in biophysical properties across major immune cell populations requires measuring many immune cell types simultaneously. Here, we take on this challenge by expanding the characterization of PM order across multiple immune cell types simultaneously. We achieve this by combining flow cytometry with a recently developed narrow-emission, plasma membrane-specific solvatochromic probe, Pro12A^39^, which enables correlative analysis of cell biophysical properties (Fig. S1). Furthermore, by sorting immune cells according to their membrane order, we establish a link between PM order remodelling and specific cell phenotype and function. Our findings demonstrate that biophysical properties represent an underexplored dimension in cell phenotyping, with the potential to enhance diagnostic capabilities and improve our understanding of cellular (dys)functional states.

## Results

### Plasma membrane order of immune cells discriminates health and disease states

To investigate the extent of PM order heterogeneity in suboptimal health conditions, we first compared the PM order in primary Peripheral Blood Mononuclear Cells (PBMCs) isolated from healthy donors (HD; 15 donors) and patients affected by chronic inflammation, namely Chronic Lymphocytic Leukemia (CLL; 5 donors) and Post-Acute Sequelae of COVID-19 infection (long COVID, LC; 15 donors) (Fig. 1A). Characterization of PM order required several key optimization steps which are described in detail in the Supporting Information (Fig. S2-S3). After optimizing protocols for sample preparation^40^ and flow cytometry settings, we maximized the sensitivity of Pro12A in detecting minute differences in PM order across a wide range of membrane compositions while ensuring its compatibility with live-cell analysis. Next, we designed a panel of cell surface markers to identify cell lineages that was compatible with our probe for PM order (i.e., minimal spectral overlap) and enabled us to identify 12 different immune cell subsets and PM order measurements using Pro12A (Fig. S4). The intensities of Pro12A, measured in two different channels, were used to calculate PM order—as Normalized Generalized Polarization (ΔGP)—for individual cells (Eq. S1-S2) where higher ΔGP values represent higher membrane order. Comparing the whole PBMC populations between healthy donors and patients (Fig. 1B-C), we observed an overall rigidification of the plasma membrane (i.e., higher ΔGP) in long COVID patients but not in those affected by CLL. However, this global effect could arise either from a uniform shift in membrane order across immune cell types or from increased membrane order in specific predominant immune cell populations. Therefore, we next analysed subtype-specific changes in plasma membrane order at the single-cell level.

**Figure 1.**
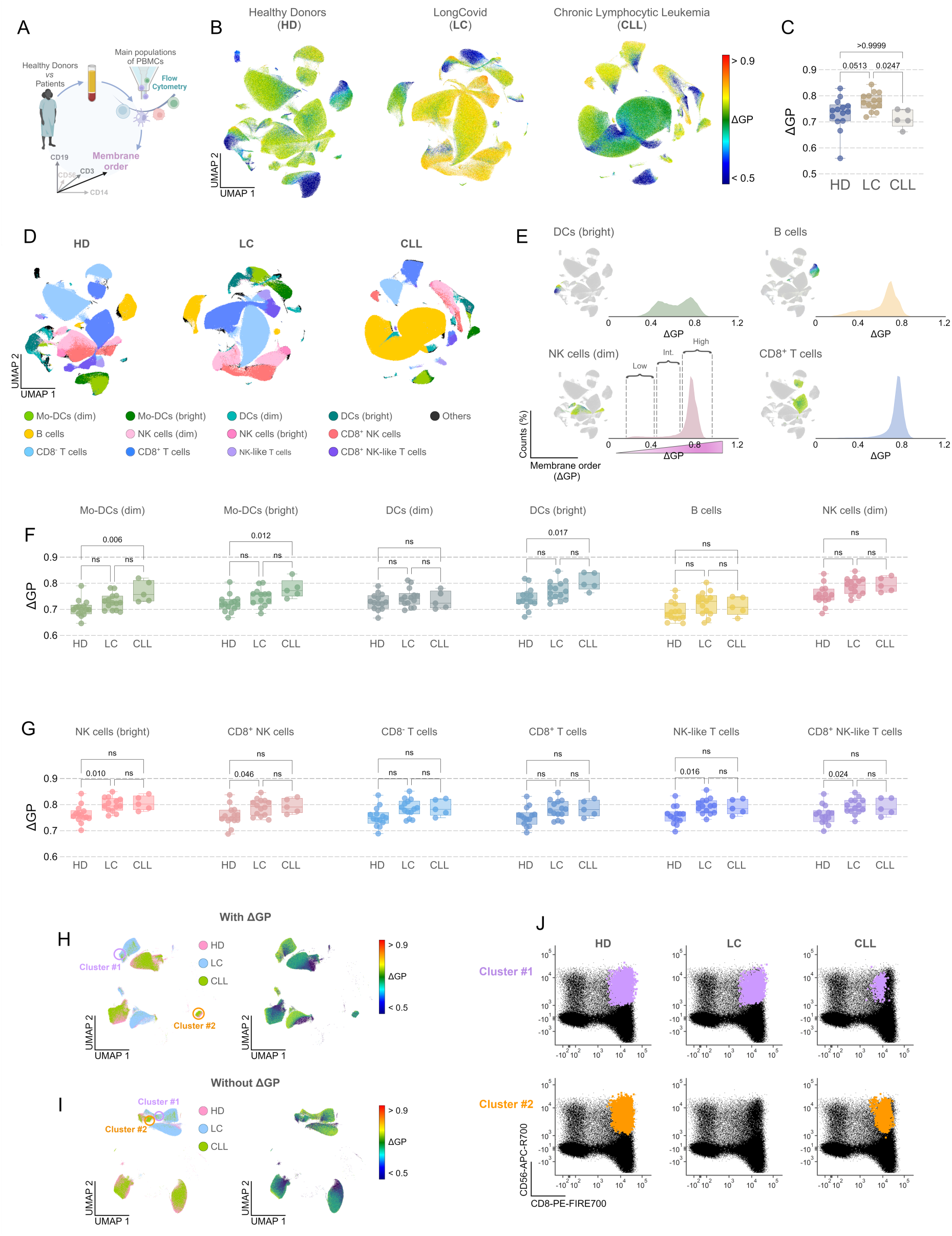
**A)** Schematic of the flow pipeline for measuring of membrane order (ΔGP) in healthy and diseased PBMCs. **B)** UMAP projections obtained from healthy and diseased PBMCs which show the distribution of membrane order values across immune cells. Pseudo colours indicate variations in ΔGP between 0.5–1. **C)** Box plots comparing the overall median ΔGP value of whole PBMCs from healthy donors and patients with long COVID and Chronic Lymphocytic Leukaemia. Each dot refers to a different donor. **D)** UMAP projections obtained from healthy and diseased PBMCs, showing the distribution of immune cell types across healthy and diseased samples. Pseudo colours indicate the different subpopulation of immune cells analysed. **E)** Pooled ΔGP distributions from selected subpopulation of immune cells from four healthy donors. The insets show the corresponding populations in the UMAP projection whereas the dashed lines mark three distinct populations of cells with different PM order. **F-G)** Box plots comparing the ΔGP values (only for high-ΔGP population) between healthy and diseased donors for a given subpopulation of immune cells:. Each dot refers to a different donor. In C, F and G, *p*-values (ns for p > 0.05) were calculated via the Kruskal-Wallis statistical test and the Dunn’s test for multiple comparison. The threshold for statistical significance was set at *p* = 0.05. Box-and-whisker plots show the median and interquartile range (25th–75th percentile), with whiskers extending to the minimum and maximum values, and all individual data points displayed. **H-I)** UMAP projections calculated from all samples–HD, LC and CLL–merged, with (**H**) or without (**I**) ΔGP as additional parameter during dimensionality reduction. The projections refer to T cell (CD3^+^) subpopulations and show the sample-specific clustering (left panel) and ΔGP values (right panel). Cluster #1 and Cluster #2 in panel H and I refer to the same gated cells within T cells subpopulation. **J)** 2D plots showing CD56 vs CD8 intensity distributions from CD3^+^ cells in Cluster #1 (purple dots) and Cluster #2 (orange dots) for each patient sample.

### Plasma membrane order is highly heterogeneous among immune cells

Using Uniform Manifold Approximation and Projection (UMAP), we clustered the 12 different immune cell subtypes by their surface markers and characterized the inter-population heterogeneity of ΔGP (Fig. 1D). A closer examination of individual immune cell subpopulations revealed different cell type specific distributions of ΔGP values. For instance, in PBMCs from healthy donors (Fig. 1E), Cytotoxic T cells and NK cells exhibited a primarily monomodal distribution, whereas Dendritic cells (DCs) and B cells followed a multimodal distribution of ΔGP values with changes in relative abundances. These findings highlight the importance of examining the distribution of ΔGP values for multiple cell types simultaneously to capture the fine details in cell type-dependent biophysical remodelling, which would be lost if we focused exclusively on descriptive parameters (means and medians) and a single cell type.

Across all cell types and samples, we observed a broad and continuous ΔGP distribution, which was categorized into three main populations: a high-ΔGP population centred at ∼0.8 ΔGP (high-ΔGP), and two smaller populations with significantly lower PM order, centred at ∼0.30 (low-ΔGP) and ∼0.55 ΔGP (intermediate, int-ΔGP), respectively (Fig. 1E, bottom-right panel exemplified for NK cells). The left-most population (low-ΔGP) exhibited ΔGP values typical of dead cells (Fig. S5A). Upon cell death, the plasma membrane becomes highly permeable, allowing Pro12A to internalize and stain intracellular organelle membranes, which have much lower order due to their specific lipid composition^9,41^.

Comparing single-donor ΔGP of the high-ΔGP population for each immune cell subpopulation, we observed general trends (Fig. 1F-G): Mo-DCs and Dendritic cells showed a systematic increase in PM order moving from healthy donors to LC and CLL patients, with CLL patients being significantly different from HD. Notably, no statistical difference was observed when comparing ΔGP values in DCs (dim) and B cells. PM order in NK and T cell subsets was slightly higher in LC and CLL patients compared to healthy donors but similar between the two patient groups. Interestingly, some cell subtypes exhibited an opposite trend for the int-ΔGP population as seen in DCs and NK cells (Fig. S5B-C).

An important observation emerged from performing dimensionality reduction on the merged datasets—i.e., HD, LC, and CLL patients analysed together—both with and without including ΔGP as an additional parameter in the UMAP analysis. For certain immune cell populations, such as NK-like cytotoxic T cells (Fig. 1H, I) and NK-like T cells (Fig. S6), incorporating ΔGP as an extra dimension provided improved resolution between subsets of cells that displayed similar expression levels of canonical markers (e.g., CD8 and CD56; Fig. 1J) but differed in ΔGP (see the increased separation between clusters #1 and #2 in Fig. 1H and 1I). Notably, cluster #2 was enriched exclusively in cells from healthy donors and CLL patients, but not from long COVID patients (Fig. 1J and Fig. S6). These results highlight the value of including PM-order phenotyping to enhance the identification and resolution of cell subpopulations.

### Cells with intermediate PM order (int-GP) represent a distinct biophysical state

While low-GP represents late apoptotic/dead cells (Fig. S5A), int-GP pool needs detailed analysis as it shows intriguing trends in diseased cells (Fig. S5). Therefore, we investigated whether the remodelling of PM order in the int-ΔGP population reflects poor cellular health or a specific cell state. First, we observed that the int-ΔGP population was primarily enriched in CD14^+^ cells (monocytes and monocyte-derived DCs) and, to a lesser extent, in B cells (Fig. S5). The enrichment was consistent across both healthy donors and patients with chronic inflammation. This suggests that certain immune cell subsets are inherently more heterogenous in PM order regardless of donor’s health status. Next, we examined cellular health in cells within the int-ΔGP gate by co-staining PBMCs from four healthy donors with fluorescently labelled Annexin-V (AnV) and a marker for cell viability. During apoptosis, phosphatidylserine (PS) accumulates in the outer leaflet of the plasma membrane^42^, leading to an overall shift in PM order and enabling Annexin-V binding^34^. However, more recently, studies have suggested broader implications of PS exposure, demonstrating its occurrence in other processes other than cell death^43–45^, even reflecting the immune cell fitness^46^.

We used two markers (AnV and cell viability) to screen this intermediate population. While AnV reports on exposed PS on the membrane outer leaflet, the viability dye interacts with amine groups overly expressed on the surface of dead cells. By using these two markers, we analysed the PM order of AnV-negative (neg-AnV, no signal for AnV and viability dye), viable AnV-positive (v-pos-AnV, positive signal for AnV but no signal for viability dye), and non-viable AnV-positive (nv-pos-AnV, positive signal for both AnV and viability dye) via flow cytometry (Fig. 2A-B, Fig. S7). We confirmed the presence of three populations, characterized by low, intermediate and high PM order that predominantly correspond to nv-pos-AnV, v-pos-AnV and neg-AnV, respectively. However, int-GP cells were present both in neg-AnV and v-pos-AnV cells (Fig. 2B, Fig. S7), indicating a pool of cells with lower membrane order within healthy cells.

**Figure 2.**
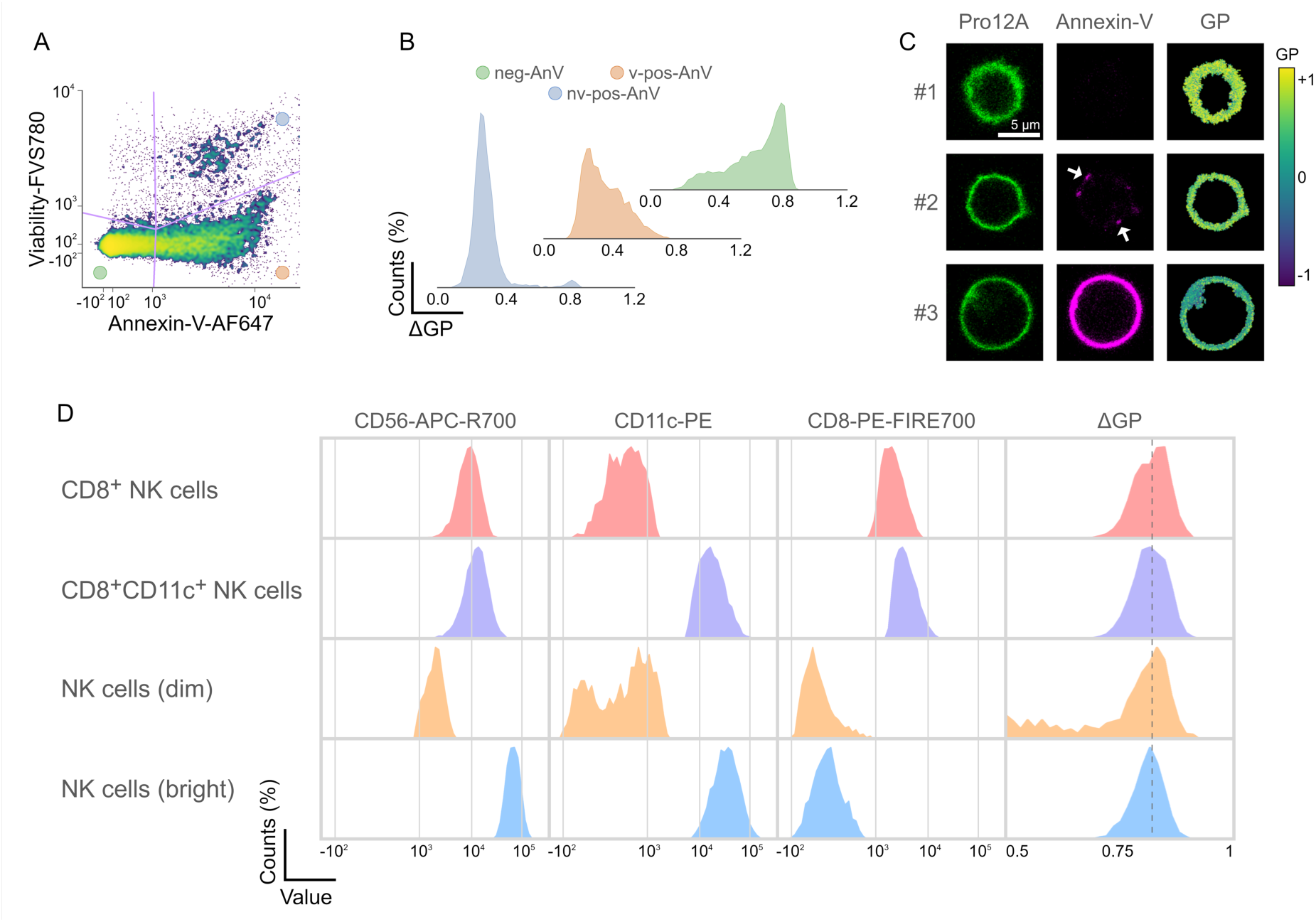
**A)** Intensity plot from cells stained with dyes for both cell viability (FVS780) and PS (Annexin-V-AF647) analysed in flow cytometry. The plot refers to T cells (CD3^+^) from healthy donors. **B)** ΔGP distributions of healthy (neg-AnV), viable AnV-positive (v-pos-AnV), and non-viable AnV-positive (nv-pos-AnV) T cells from one of four healthy donors. **C)** Microscopy images acquired with a spectral confocal microscope, showing three different cells (#1, #2 and #3) after being stained with both Pro12A and Annexin-V-AF647. Green, magenta and pseudo-coloured images refer to Pro12A, AnV and calculated GP values, respectively. The scale bar indicates 5 µm distance and the white arrows indicate AnV puncta. **D)** Histograms showing the intensity distribution of different biomarkers and the distribution of ΔGP values (far right column) for each subpopulation of NK cells from long COVID patients.

To assess whether the decrease in ΔGP was due to a reduction in PM order or partial internalization of Pro12A, we used spectral confocal microscopy which supported our flow cytometry observations (Fig. 2C) and confirmed the existence of three distinct cell populations and found no evidence of Pro12A internalization in cells with int-ΔGP. These cells displayed a punctate accumulation of AnV. A similar pattern has been reported in T cells upon receptor-mediated activation and attributed to localized membrane scrambling leading to PS exposure^44^ and externalized PS on T cells is suggested to be an extrinsic inhibitory molecule in cellular exhaustion^46^.

When we profiled lipid order along the membrane, we observed a global reduction in ΔGP, not just at the AnV clustering sites (Fig. S8). This suggests that membrane order remodelling is more global than localized to PS^+^ spots which could be caused by partial breakage of membrane asymmetry^47^ but not the membrane integrity. In addition, cells with lower ΔGP showed a slight decrease in the main surface markers for cell lineage (i.e., CD3, CD19, and CD56), suggesting that these cells might be a different subset (Fig. S9). Altogether, our findings shows that int-ΔGP pool is not apoptotic cells (tested with AnV) and instead corresponds to viable cells with a different biophysical state than high-ΔGP pool, positioning PM order as a potential new dimension in immune cell phenotyping.

### Plasma membrane order is an orthogonal parameter to canonical cell surface markers

After exploring that ΔGP can be an indication of health state (Fig. 1) and high-ΔGP and int-ΔGP pools might represent different cell states (Fig 2A-C), we asked whether ΔGP can be correlated with surface biomarkers, which are commonly used to identify cell types, states and diseases. Notably, we found orthogonality between ΔGP and most cell biomarkers, with their correlation (either positive or negative) depending on the cell type and sample under examination (Fig. 2D, Fig. S10–S13). For instance, when we compared PM order in CD8⁺ NK cells, we observed that cells with higher PM order have the same level of CD56 and CD8 but lower CD11c signal, suggesting a negative correlation between ΔGP and CD11c (Fig. 2D, upper two rows and Fig. S13). In contrast, for NK cells with low CD56 expression (NK cells (dim)), lower CD11c expressing cells showed lower ΔGP (hence a positive correlation in this case between ΔGP and CD11c). In comparison, NK cells with high CD56 expression (NK cells (bright)) showed higher CD11c expression and higher ΔGP values (Fig. 2D, lower two rows, Fig. S13). These findings show that PM order might correlate with protein expression but not universally. Therefore, PM order remodelling may be associated with different dynamic cell states and could potentially define immune cell subsets with complex surface signature (as seen in Fig. 2D with three differentially expressed markers on the surface). Considering the complexity of the surface marker signature and its correlation with cellular fitness; PM order will be an easy and straightforward way to access functionally different cell states. Therefore, we next performed sorting based on PM order and assessed the functionality of different pools.

### Biophysical sorting based on PM order

Having observed the link between PM order and the expression of surface biomarkers, we next aimed to test whether ΔGP could be used to isolate immune cells with distinct functional states and properties. For this, we first established the sorting of NK cells solely based on their PM order and performed functional assays on NK cell subsets characterized by either low (NK-L) or high (NK-H) PM order. We began by enriching NK cells from whole PBMCs isolated from multiple healthy donors and performed biophysical sorting of these cells. Specifically, we collected cells whose PM order fell below the 15th percentile (NK-L) and above the 85th percentile (NK-H) of the total ΔGP distribution after excluding dead cells (Fig. 3A, Methods for details). Sorted cells were either analysed immediately or cultured in recovery medium for 24 hours prior to functional assays.

**Figure 3.**
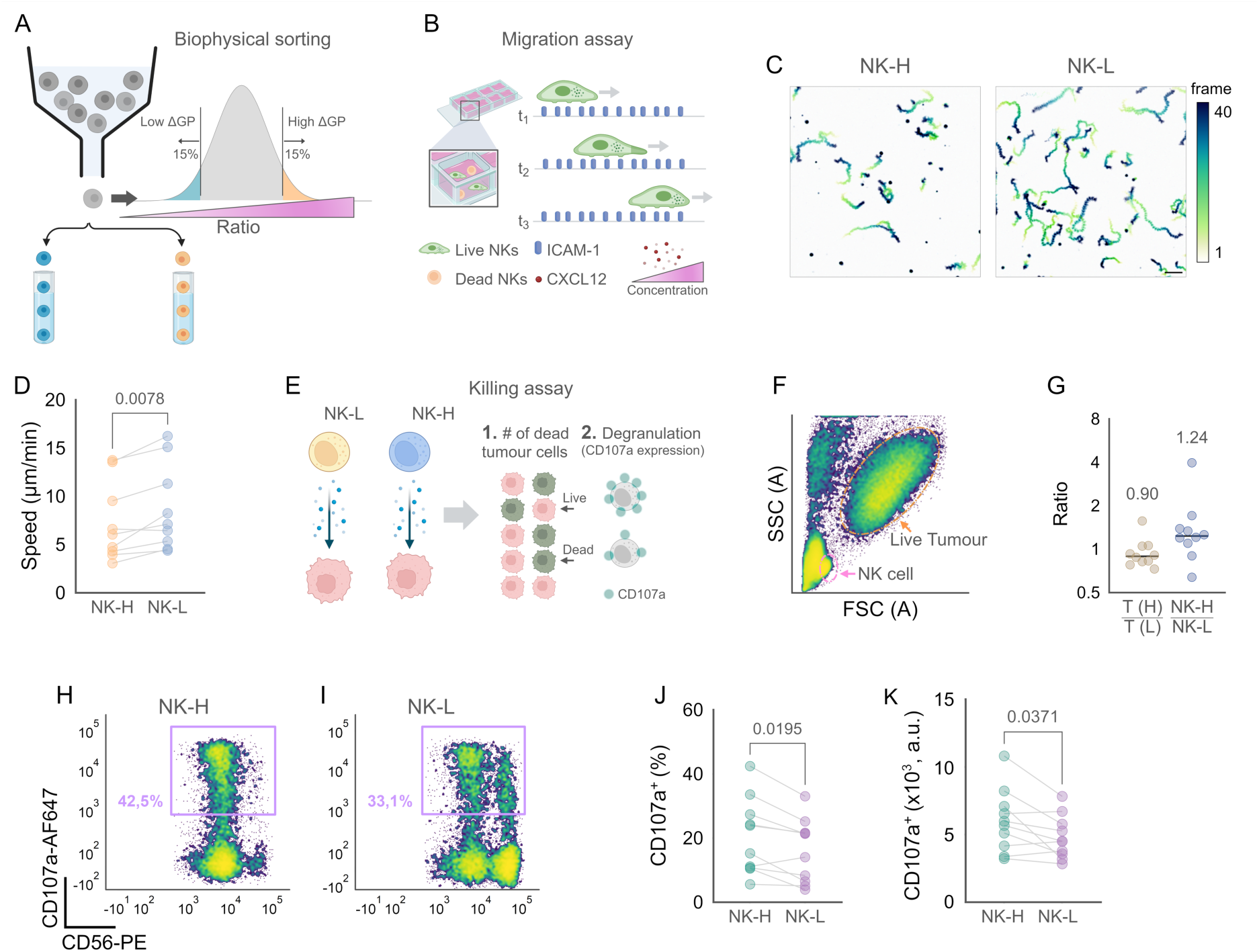
**A)** Schematic of the NK cell sorting based on their PM order. **B)** Schematic of the migration assay performed on NK-H and NK-L subsets. **C)** Example of migration trajectories recorded over a 40-frame period using confocal microscopy for the two NK cell subsets. **D)** Mean migration speed of the two NK cell subsets from individual donors (n = 9). **E)** Schematic of the killing assay performed on NK-H and NK-L subsets. **F)** Forward versus side scatter plot from flow cytometry analysis of NK cells after incubation with K562 tumour cells. **G)** Ratios between number of live tumour cells from the high PM order-fraction (H) and low PM order-fraction (L), and between number of cells from NK-H and NK-L subsets, from individual donors (n = 10) after incubation with K562 tumour cells. The numbers refer to the mean ratio value. **H-I)** CD107a versus CD56 intensity plots from flow cytometry, showing the fraction of NK-H and NK-L cells that underwent degranulation upon incubation with K562 tumour cells. **J-K)** Percentage (J) of CD107a-expressing NK cells and mean CD107a intensity (K) in the two NK cell subsets from individual donors (n = 10). Statistical analysis in D, J and K plots was performed via Wilcoxon matched-pairs signed rank test.

### Lower PM order correlates with faster migration of NK cells

The efficiency of NK cells depends on their ability to migrate towards the target cells and physically engage to induce cell death. Therefore, we tested both the migration and the killing efficiency of sorted NK cells as a function of their PM order. For the chemotaxis assays to assess potential differences in migration, NK-L and NK-H cells were seeded on ICAM-1–coated coverslips and exposed to a CXCL12 chemokine gradient (Fig. 3B). Analysis of migration trajectories (Fig. 3C) revealed that NK-L cells —characterized by a more fluid plasma membrane— migrated significantly faster than NK-H cells (Fig. 3D). Despite high inter-donor variability, each donor exhibited a consistent trend, with NK-L cells migrating approximately 25% faster on average. Several mechanisms may underlie this observation, since PM order can directly or indirectly influence cell migration at multiple levels. To reveal whether the PM order is directly involved in migration capacity, we measured the PM order after recovery and observed that the difference in ΔGP between the two populations normalized after 24h of recovery in culture conditions, converging to an intermediate value (Fig. S14). This suggests that PM order is an indicator of the activity at the time of selection but not a direct determinant of the migration of NK cells at the time of measurements, which will be discussed later.

### NK cells with increased PM order show higher killing efficiency

We continued our functional characterization of biophysically sorted NK cells by performing killing assays (Fig. 3E). After 24 hours of recovery in cell culture medium, NK-H and NK-L cells were each mixed with an equal number of K562 tumour cells and incubated for 6 hours. Subsequently, cells were stained with a cocktail of antibodies, including a viability dye and CD107a—a marker of degranulation of NK cells—and analysed by flow cytometry. The cytotoxic efficacy of NK cells was assessed using two independent approaches: *(i)* by calculating the ratio of live NK-H cells to NK-L cells (NK-H/NK-L) and the ratio between live tumour cells from the two subsets (i.e., T(H)/T(L)), and *(ii)* by measuring CD107a expression levels. CD107a upregulation at the cell surface indicates the extent of degranulation^48^ while the ratio of live NK-H/NK-L cells and T(H)/T(L) reflect the effectiveness of degranulation and NK cell resilience to self-killing. Live tumour cells were identified based on light scatter characteristics (Fig. 3F), while live NK cells were gated as CD56-positive and viability dye-negative (Fig. S14). Looking at the ratios of live tumour cells and live NK cells between the NK-H and NK-L subsets (Fig. 3G), cells with higher PM order killed ∼10% more tumour cells and maintained approximately 24% higher viability compared to NK-L cells. The moderate increase in number of killed targets could be because sorted NK cells did not undergo any interleukin-based activation prior to incubation with tumours. Furthermore, NK cells with higher PM order showed a 28% increase in the frequency of degranulating cells (Fig. 3H-J, Fig. S14) and a 20% higher CD107a signal intensity on average (Fig. 3K, Fig. S14). Collectively, our results suggest that while the actual killing efficiency was moderately better in our experimental setup, NK-H cells can potentially be better killers due to both enhanced survival after degranulation and more effective release of lytic granules. Since we previously observed that PM order converges in NK-L and NK-H pools during recovery in culture conditions, it is likely that we selected different functional cell subtypes or states by biophysical sorting; and PM order is an indicator of cell states but not a direct determinant for killing. For example, regardless of exposure to tumour cells, NK-H were predominantly CD56^dim^, whereas NK-L also contained a detectable population of CD56^bright^ cells (Fig. S14). For a detailed molecular identification of the cells states of NK-L vs. NK-H pools, we next performed transcriptomics analysis of sorted pools.

### Higher PM order in NK cells is associated with reduced immunosuppression

When we analysed the transcriptome of NK-L and NK-H, 261 genes were significantly downregulated in NK-H compared to NK-L cells, while 37 genes were significantly upregulated (Fig. S15 and Supplementary Table on genes list). The most significantly upregulated genes including RND1, NEU1, DHCR7, ZNF503-AS1 and SLC25A4 (Fig. 4A), play a role in cellular stress response and membrane dynamics^49–52^. On the other hand, the most significantly downregulated genes–MYBL2, TYMS, STMN1 and TK1–are associated with cell cycle regulation and proliferation^53–57^. Gene Ontology (GO) annotation using Gene Set Enrichment Analysis (GSEA), revealed a significant enrichment of downregulated genes that were localized to the external plasma membrane and the membrane of secretory granules (Fig. 4B). In terms of molecular function, NK-L cells showed significant downregulation of genes in pathways associated with immune receptor activity, particularly cytokine receptors (Fig. 4C). In the biological process domain, significantly downregulated transcripts in NK-H were enriched in pathways involved in cell cycle regulation, interleukin production, cytokine signalling, and chemotaxis (Fig. 4D).

**Figure 4.**
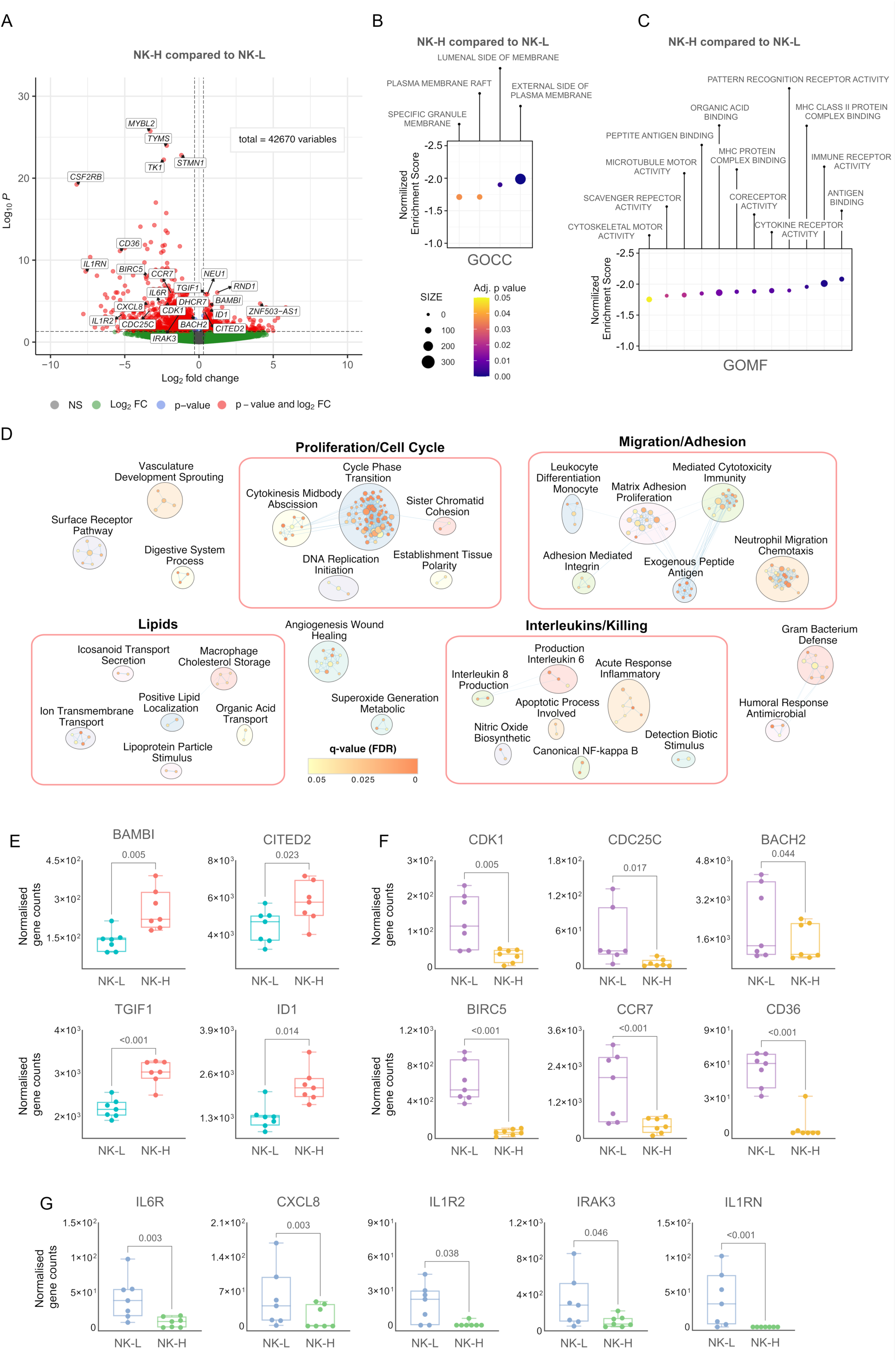
**A)** Volcano plot showing the differential gene expression in the NK-H subset compared to NK-L. **B-D)** Results from Gene-Set Enrichment Analysis (GSEA) based downregulated gene ontology (GO) annotations within the cellular components (B), molecular function (C) and biological process (D) domains in the NK-H subset compared to NK-L. **E-G)** Comparison of normalized gene counts between NK-L and NK-H subsets for selected genes from different donors (n = 7). In panel B,C and D, the adj. p-values, are calculated using a permutation-based approach followed by correction for multiple comparisons using the false discovery rate (FDR) method, while in panels A, E-G the adjusted p-values are calculated using a generalized linear model (GLM) and corrected for multiple comparison using the Benjamini-Hochberg (BH) method. Differences were considered significant for adj. p-values ≤ 0.05. Box-and-whisker plots show the median and interquartile range (25th–75th percentile), with whiskers extending to the minimum and maximum values, and all individual data points displayed.

Based on these pathway-level changes, we further examined individual genes that were significantly different between NK-H and NK-L cells. We first analysed whether the differences in membrane order would affect the expression of activating or inhibitory surface receptors. This is relevant because NK cell cytotoxicity against K562 tumour targets is mediated by receptor–ligand interactions, involving, for instance, NKG2D overexpression on NK cells^58^ or upregulation of DNAM-1 ligands or TRAIL receptors (DR4/DR5) on K562 cells^59,60^. Although some genes related to receptor-mediated cytotoxicity were differentially expressed, the differences between NK-H and NK-L subsets were not statistically significant (Supplementary Table on genes list). Interestingly, several differentially regulated genes were enriched in the Transforming Growth Factor-beta (TGF-β) signalling pathway from KEGG (Fig. S16). TGF-β is a multifunctional cytokine and a key mediator of immunosuppression^61^. In NK cells, TGF-β promotes the conversion of cytotoxic NK cells into less cytolytic ILC1-like cells^62^, inhibits activation and cytotoxicity via repression of the mTOR pathway^63^, reduces IFN-γ production, and downregulates activating receptors such as NKG2D and NKp30^62^. A recent study demonstrated that knockout of SMAD4, a downstream mediator of TGF-β signalling, preserves NK cell cytotoxicity^64^. In our transcriptomic analysis, we identified four genes that were significantly upregulated in NK-H cells compared to those with lower membrane order (Fig. 4E): BAMBI and TGIF1, both negative regulators of TGF-β signalling^65,66^, and CITED2 and ID1, which are context-dependent modulators of TGF-β^67–72^. These findings suggest that the enhanced killing capacity of NK-H cells might be linked to a functional inhibition of the crucial pathways including TGF-β pathway.

### NK-L cells exhibit a more proliferative but less cytotoxic phenotype

While NK cells with increased PM order show indications of reduced TGF-β-dependent signalling, the NK-L subset displayed a genetic signature suggestive of increased proliferative potential. For example, NK-L cells showed significant upregulation of genes encoding Cyclin-dependent kinase 1 (CDK1) and Cell Division Cycle 25C (CDC25C, Fig. 4F). Their gene products regulate the transition from late G2 phase into mitosis and are mechanistically interconnected promoting a more proliferative state^73^. Given the enhanced proliferative phenotype, we assessed whether NK-L cells might be undergoing a form of replication-induced exhaustion. However, our transcriptomic analysis did not reveal significant changes in the expression of exhaustion-associated inhibitory markers such as TIGIT, LAG-3, KLRC1, and PDCD1^74^, nor in activating markers such as FCGR3A and KLRK1 (NKG2D)^75^. Conversely, NK-L cells showed significantly increased expression of BACH2 and BIRC5 (Survivin, Fig. 4F). BACH2 is a transcription factor and intrinsic negative regulator of NK cell maturation and cytotoxicity, typically overexpressed in immature CD56^bright^ NK cells. Its regulation has been linked to Granzyme B (GzmB) production^76^. Similarly, BIRC5, commonly upregulated in proliferating cells^77^, has been associated with reduced levels of intracellular cytotoxic proteins such as perforin, GzmB, TNF-α, and IFN-γ^78^. In addition, NK-L cells significantly overexpressed CCR7 and CD36 (Fig. 4F). CCR7 is preferentially expressed on immature CD56^bright^ NK cells, mediating homing to secondary lymphoid organs^79^. On the other hand, CD36 is responsible for fatty acid uptake and it has been shown to drive β-oxidation in cells upon infection^80^. Interestingly, impairment of fatty acid metabolism via inhibition of β-oxidation correlates with decreased capacity of NK cell migration during acute retroviral infection^81^ linking higher CD36 levels to enhanced migration.

### NK-H cells show an interleukins-dependent pro-inflammatory phenotype

Notably, the NK-H subset showed a significant downregulation of several genes associated with interleukins (Fig. 4G). For instance, we observed transcriptional downregulation of CXCL8 (IL-8) and IL6R. The latter encodes for the Interleukin-6 (IL-6) receptor, and IL-6 has been shown to downregulate cytotoxicity of NK cells via reduction of perforin and granzyme B levels^82^. Remarkably, the NK-H subset showed significant downregulation of three different genes–IL1R2, IL1RN and IRAK3–which are involved in the suppression of the Interleukin-1 (IL-1) dependent inflammatory response. Therefore, downregulation of IL1R2, IL1RN and IRAK3 indicate that NK cells with higher PM order might represent a subset of cells more sensitive to IL-1-mediated inflammation.

### PM order correlates with maturation and cytotoxicity in NK cells

Although the membrane order of both subsets equalized to an average value after 24 hours of resting, the two groups presented distinct killing and migratory capacity in our post-resting functional assays (Fig. 3). While transcriptomic analysis revealed phenotypic differences in RNA level between freshly sorted NK-L and NK-H subsets, we asked whether PM order might be representative of a more constitutive phenotype of cells which is retained even after the culture conditions. For this, we focused on the cell surface architecture as it predominantly determines the cellular activity. Thus, we performed the proximity network assay (PNA) to assess the surface protein interactome of single NK cells^83^. This assay uses 155 antibodies against immune cell surface proteins functionalized with DNA barcodes and provides information of protein abundance, clustering and colocalization with other proteins, generating a 3D spatial network of surface proteins for each single cell (see Methods for details). Compared to transcriptomics, PNA is a direct measure of surface proteins; and compared to proteomics, PNA shows the surface specific molecules in addition to their clustering and proximity to each other. Using PNA, we measured the abundance and physical redistribution of selected NK markers at the cell surface of NK cells isolated from healthy donors and sorted based on PM order (i.e., NK-H and NK-L). UMAP projection from PNA analysis of pooled donors (Fig. 5A) showed the presence of three main clusters which were differently enriched in NK-H and NK-L subsets (Fig. 5B): Cluster 1, described by a phenotype typical of more mature NK cells (i.e., high expression of CD16, CD57, KIRs); Cluster 3 characterized by high expression of CD56, NKG2A, CD27 and CD94 among the others, indicating immature NK cells^84^; and Cluster 2, showing a phenotype intermediate between Cluster 1 and 3 (Fig. 5C). Cluster 3 had also higher expression of NKG2C marker associated with adaptive-like NK phenotype^85^. Notably, Cluster 1 (more mature NK cells) was enriched in NK-H cells whereas NK-L consisted of less Cluster 1 and significant more Clusters 2 and 3 (Fig. 5B and S17A), confirming observations on NK cells subsets during killing assays. Besides exhibiting a more mature phenotype, NK-H cells displayed a distinctly cytotoxic profile, with elevated expression of CD16, CX3CR1, KLRG1, and Siglec-7/9—markers characteristic of terminally differentiated CD56^dim^ NK cells with heightened effector function (Fig. 5D and S17B). In contrast, NK-L cells retained higher levels of HLA-DR/DP/DQ, HLA-ABC, CD58, and CD48, features associated with activation, adhesion, and antigen-presentation–like states. These differences likely account for the reduced killing efficiency observed in our functional assay, as CD56^bright^-like NK cells (NK-L) are predominantly cytokine-responsive rather than cytotoxic, whereas CD16^high^ NK-H cells exhibit enhanced cytotoxic potential^86^.

**Figure 5.**
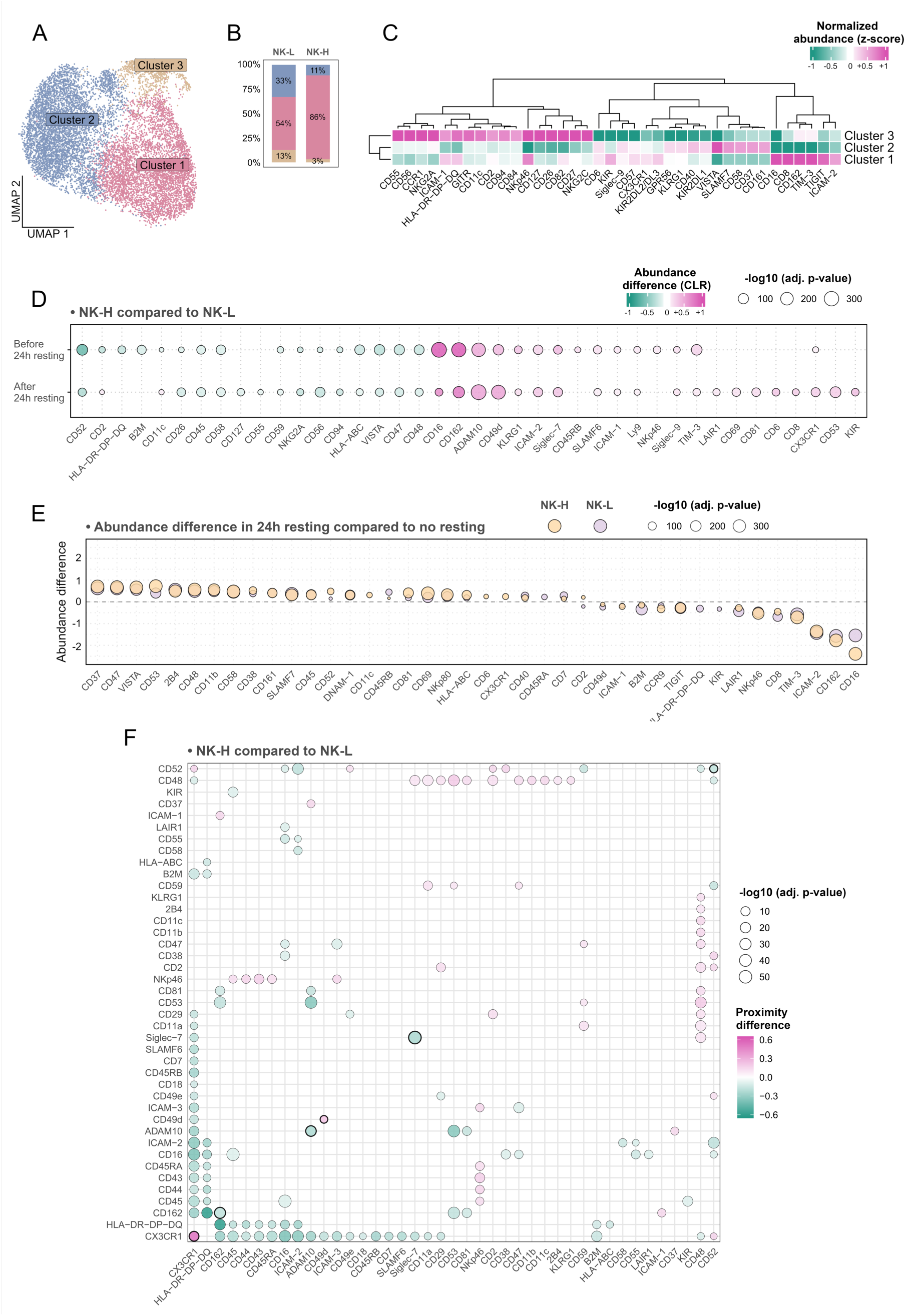
**A)** UMAP projection from PNA analysis of sorted NK cells from different healthy donors (n = 2) pooled together, showing the presence of three distinct clusters (Cluster 1, Cluster 2, and Cluster 3). **B)** Box plots showing the percentage composition of NK-H and NK-L in clusters 1, 2, and 3 before 24 resting. **C)** Heatmap from PNA analysis showing differential abundance (z-score) of selected surface markers among the three NK-cell clusters. **D–E)** Differential abundance of surface markers from PNA analysis, showing changes occurring after 24-hour recovery in culture (D) or intrinsic phenotypic differences between NK-H and NK-L subsets (E). **F)** Differential Proximity Score plot comparing proximity scores of surface markers in CD56^dim^CD16^bright^ cells from NK-H versus NK-L subsets. Magenta dots denote markers with higher proximity scores in the NK-H subset.

### Phenotypical differences between sorted NK-H and NK-L are preserved after resting

One of the key questions from our previous data (Fig. 3) was why NK-H and NK-L cells displayed distinct functional behaviors despite equalization of PM order. To investigate this, we compared the PNA analysis before and after a 24-hour recovery period. This experiment showed that culture-induced remodeling affected both subsets similarly (Fig. 5E and Fig. S17C). In both NK-H and NK-L cells, resting led to increased expression of early activation markers such as CD69 and CD38 (Fig. 5E)^87^. Activating receptors—including DNAM-1, 2B4, SLAMF7, and NKp80—also increased, consistent with enhanced target recognition and cytotoxic signalling potential^88^. Adhesion and synapse-associated molecules (CD58, CD48, CD11b, and the tetraspanins CD37/CD53/CD81) likewise increased, reflecting a shift toward a more synapse-competent phenotype driven by in-vitro culture conditions. Meanwhile, decreases in NKp46, TIGIT, TIM-3, and LAIR-1 likely reflect activation-dependent receptor modulation. Reduced CD16 expression has also been widely reported in cultured NK cells as a consequence of ADAM-mediated shedding following activation^89^.

### Biophysical sorting based on PM order can further dissect heterogeneity within NK cell populations

Notably, both NK-H and NK-L subsets contained CD56^dim^CD16^pos^ NK cells (Figure 5B), a population classically defined as more mature and cytotoxic (Figure S17). Despite sharing this phenotype, the two subsets displayed strikingly different PM order, prompting us to examine differences in surface protein abundance and spatial distribution via PNA (Fig. S17D, E) ^90^. Notably, the two subsets exhibited clear differences in several key markers, suggesting that CD56^dim^CD16^pos^ NK-H and CD56^dim^CD16^pos^ NK-L cells have nuanced differences in their surface state. Specifically, CD56^dim^CD16^bright^ NK-H cells expressed lower levels of CD48, CD52, and CD47, molecules typically involved in inhibitory signalling, suggesting that they can be more primed for activation. Moreover, the NK-H subset displayed higher levels CD162 (PSGL-1) and CD49d. These changes suggest that there might be diverse functional subsets of NK cells exhibiting similar canonical marker expression, but different overall membrane order and surface proteome. To dissect this better, we next examined the surface protein network on NK-H and NK-L cells next using the PNA analysis.

### NK-H and NK-L cells have different surface protein network

PNA, beyond the abundance data, yields a colocalization matrix of all the tested proteins. When we examined this spatial proximity map of surface proteins in NK-H cells compared to NK-L cells (i.e., differential proximity map, Fig. 5F), we observed several interesting candidates exhibiting a change in their proximity. For example, we observed that higher membrane order correlated with reduced proximity scores—*i.e.,* less colocalization—between ADAM10 and the major tetraspanins CD37, CD53, and CD81. Although published work shows that ADAM10 interacts directly only with CD9 and members of the TspanC8 subgroup^91,92^, our findings suggest a broader relationship between tetraspanins and ADAM10 which is shown to have large impact on surface accessibility^93^. We also detected substantially decreased proximity between CD53 and CD162 (a substrate for ADAM10) and increased proximity between CD53–CD48. Importantly, except for CD53, no differences in tetraspanin abundance were detected between CD56^dim^CD16^bright^ NK-H and NK-L subsets (Fig. S17D), yet their spatial organization differed markedly. This supports the idea that PM order, independent of expression level, can be an indicator of NK-cell functional potential by correlating with the complex protein network at the cell surface.

## Discussion

Remodelling of PM order—one of the key biophysical properties of living cells—has been demonstrated to be important for the overall immune system fitness. Here, we present a new approach that integrates multiparametric flow cytometry with biophysical sorting to quantify PM order and link it to specific functional states. By combining lineage-defining markers with environment-sensitive solvatochromic dyes, our workflow enables high-dimensional phenotyping of thousands of PBMC-derived cells at single-cell resolution, providing access towards the full heterogeneity of membrane biophysical states across the circulating immune system.

Using this strategy, we observed striking variability in PM order across immune subsets from healthy donors, patients with chronic lymphocytic leukemia (CLL) and individuals with long COVID (LC). Disease states were characterized by a consistent shift toward increased membrane order in multiple immune cell types, potentially reflecting the lipid metabolic dysregulation and dyslipidaemia reported in these conditions^94,95^. These results raise the possibility that disease-specific or cell type-specific PM-order signatures may emerge depending on the nature of the immune insult which requires further investigation^96–101^. Importantly, the single-cell resolution of our approach allowed us to detect nuanced, subset-specific shifts in PM order that would be obscured by bulk readouts, suggesting that distinct immune lineages remodel their membranes differently even under shared systemic perturbations. Correlations between PM order and additional phenotypic markers further indicate that membrane order can act as an orthogonal reporter of transient cellular states.

We then applied biophysical sorting to NK cells and performed functional, transcriptional and proximity network characterization of subsets separated solely by their PM-order profiles. This analysis revealed distinct NK-cell states differing in cytotoxic potential, migratory capacity, transcriptional programs and surface protein network. Although culture conditions led to homogenization of PM order across pools, functional and phenotypical differences persisted, implying that PM order reflects deeper regulatory states rather than transient physical properties alone. This was confirmed with PNA analysis before and after the 24h resting. These findings highlight membrane order as an informative dimension that captures nuanced immune-cell states and function.

More broadly, our results support the concept that PM order is an adaptive biophysical variable shaped by both intrinsic programs and extrinsic cues. Within diseased tissues, PM-order remodelling may arise from receptor activation, metabolic rewiring, lipid imbalance or mechanical stress. Given that membrane order influences processes such as immune synapse stability, migration and effector responses^12,102^, quantitative assessment of PM order provides a straightforward means to detect pathological immune states as a proxy of complex molecular remodelling. Solvatochromic probes thus offer a powerful chemical-biology toolkit to interrogate inflammation arising from infection, tumorigenesis, metabolic dysfunction or dietary factors.

Finally, our cytometry-based platform is compatible with standard commercial instrumentation and can be extended to additional biophysical dimensions through organelle-targeted and mechanosensitive probes^103^. As interest in biophysical determinants of immune function continues to grow, this work establishes PM order as an orthogonal and broadly applicable parameter, with potential for integration into routine immunophenotyping, systems immunology and diagnostic workflows.

## Methods

### Synthesis of BSLBs with different membrane order

Silica beads and lipid species (Cholesterol, 1,2-dioleoyl-sn-glycero-3-phosphocholine (DOPC), 1-palmitoyl-2-oleoyl-glycero-3-phosphocholine (POPC) and 1,2-dipalmitoyl-sn-glycero-3-phosphocholine (DPPC)) were purchased from Bangs Laboratories and Avanti Polar Lipids, respectively. Synthesis of BSLBs consisted of two steps: *i)* formation of liposomes with given PM order and *ii)* coating of silica beads with a specific liposome composition. *Formation of liposomes*. For each lipid species, 10 µL of lipid solution in Chloroform (25 mg/mL) was transferred to a 1.5 mL Eppendorf and dried out under a Nitrogen flow. Then, 1 mL of *ad-hoc* buffer (SLB buffer: 10 mM HEPES, 150 mM NaCl, pH=7.4) was added to hydrate the lipid layer for 5 minutes under vigorous vortexing, and the solution was sonicated for 10 minutes (tip sonication, power=3, duty cycle=40%) to yield ∼30 nm-sized liposomes. *Coating of silica beads*. 5 µm-sized beads were washed 2x with SLB buffer, resuspended in 45 µL of liposomes solution and diluted to 500 µL final volume in SLB buffer. Complete coating of silica beads was achieved after 30 minutes in bath sonication, after which beads were washed 2x in SLB buffer to remove excess of liposomes and stored a +4 °C.

### Optimising resolution for PM order measurements

In our study we used Pro12A to probe PM order (Fig. S1)^39^ both in synthetic and biological membranes. We prepared BLSBs with either low (DPPC-Cholesterol 1:1 coated beads) or high (DOPC coated beads) membrane order and used them to identify the best PMT voltages in the blue and green channels of the flow cytometer (BD LSRFortessa). 50 µL of liposomes-coated beads was diluted into 500 µL of SLB buffer and stained with 1.5 µL of Pro12A under vortexing. Acquisition of flow data started after 1 minute from dye addition. From the theoretical map of ΔGPs *vs* amplification factors (Fig. S2), we selected two different areas of low and high intensity signals and performed a voltration in one channel while keeping the PMT in the other channel constant. By doing so, we were able to identify the optimal pair of PMT voltages which ensured the highest resolution (*i.e.*, largest ΔGP between low and high order membranes) and reproducibility of GP measurements. For each lipid composition, we prepared four independent batches of BSLBs and acquired ≥ 5×10^3^ beads per measurement to ensure statistical robustness.

### Pro12A internalization and cytotoxicity

We assessed degree of internalization and cytotoxicity of Pro12A by acquiring images in spectral confocal microscopy (LSM780 Zeiss microscope) over time and at different concentrations. Dye was considered internalized and cytotoxic when signal from the intracellular environment became visible or blebbing of the plasma membrane started occurring, respectively (Fig. S3). Internalization of Pro12A was studied using CEM/C1 cells which were cultured in RPMI-1640 medium supplemented with 10% foetal bovine serum (FBS, Sigma Aldrich) at 37 °C and 5% CO2. 8-wells ibidi chambers were precoated with BSA (3 mg/mL) for 30 minutes and 100 µL of CEM/C1 cells (6×10^6^ cells/mL in Leibovizt (L-15) medium) were added to each well and topped up with additional 300 µL of fresh L-15 medium. To each well, 1 µL of Pro12A was added to a final concentration of either 0.25 µM or 2.25 µM, and t-stack images were acquired (10 images every 90 seconds). Cell-specific plasma membrane order was calculated using our previously released software^104^.

### Study Participants

The present study analysed PBMCs from 13 healthy donors, 13 Long COVID patients and 5 CLL patients. For healthy donors, cells were isolated either from anonymous Buffy Coats (7 donors) or blood samples (6 donors raging from ages 27 to 44 years). Long COVID patients were characterized with >12 weeks of objectively measurable organ damage/dysfunction following PCR-verified SARS-CoV-2 infection and not explained by the severity of acute COVID or treatments for the same. The analysis included 12 post-acute sequelae of COVID-19 (PASC) patients and 1 recovered PASC patients (PASC-R). Long COVID patients ranged from ages 35 to 64 years. Venous blood samples were collected at a single time point for each individual. Informed consent was obtained from all patients. The Swedish ethical permit is covered under Dnr 2021-03293. Samples from Chronic Lymphocytes Leukaemia (CLL) patients were collected after 3 months from the second dose of vaccination against COVID-19. Patients were in their early-stage untreated CLL.

### PBMCs isolation

PBMCs were isolated either from Buffy Coat (healthy donors) or blood (Long COVID and CLL patients) following a different protocol, depending on the sample. healthy donors and CLL patients: 45 mL of buffy coat was diluted 1:1 with PBS into 2×50 mL Falcon tubes and properly mixed. 30 mL of diluted solution was transferred in 2 new Falcon tubes pre-filled with 15 mL of Ficoll, using gravity-driven flow to overlay the diluted blood onto Filcoll. The tubes were centrifuged at 400 g for 25 minutes (brake off). The fraction containing PBMCs was collected into 2 new Falcon tubes and washed 2x with 45 mL PBS. During the washing steps, cells were pelleted by spinning the tubes at 300 g for 5 minutes (brake on). After the last washing, cells were resuspended in 20 mL PBS, filtered through a cell strainer, and pelleted one last time at 200 g for 10 minutes. After removal of the supernatant, cells were resuspended at a density of 10^8^ cells/mL in a solution for cryopreservation (in 90% FBS + 10% DMSO) and stored in 1 mL batches at −140 °C until the analysis. *Long COVID patients:* Following plasma collection, blood was processed to isolate peripheral blood mononuclear cells (PBMCs) as follows: briefly, Lymphoprep™ (STEMCELL Technologies) was first added to SepMateTM tubes (STEMCELL Technologies) by carefully pipetting through the central hole of the SepMate insert, after which pre-diluted blood (Blood mixed with PBS + 2% FBS 1:1) was layered on the top of Lymphoprep™ (1 part of Lymphoprep: 2 parts of diluted blood) by pipetting it down the side of the tube carefully by minimizing the mixing of blood with Lymphoprep™ and centrifuged at 1200 x g for 10 minutes at ambient temperature (15 - 25°C) with the brake on. The top layer containing the enriched mononuclear cells (MNCs) was poured off into a new 50 mL tube and washed twice with PBS+2% FBS by centrifuging at 300 x g for 8 minutes at ambient temperature, with the brake on. Cells were then counted and cryopreserved (in 90% FBS + 10% DMSO) at a concentration of 5 million PBMCs/ml per cryotube stored at −80 °C first followed by transferring them to a liquid nitrogen tank for long-term storage.

### Immune staining

Frozen PBMCs were thawed by adding a 10-fold volume of RPMI-1640 (Sigma-Aldrich) supplemented with 10% FBS (Sigma-Aldrich) and Benzonase Nuclease (10 units/mL, Sigma-Aldrich), pelleted at 400 g for 8 minutes, resuspended in 5 mL medium and allowed to recover. Since previous studies have shown that post-thaw resting of PBMCs may or may not be beneficial depending on the cell type^105^, we assessed the effect of different resting times after thawing on the PM order of major immune cell populations and compared the results with freshly isolated PBMCs analysed immediately after isolation (Fig. S18). On average, we observed a 6–12% decrease in PM order following freeze–thawing, with no significant differences between 1, 4, or 8 hours of recovery. However, prolonged incubation in culture medium led to an increase in apoptotic cells, particularly among monocyte-derived dendritic cells. Based on these findings, we opted for a minimum 1-hour recovery period—sufficient to restore membrane biophysical properties while minimizing apoptosis and ensure complete recovery of Monocytes and Dendritic cells. After recovery, PBMCs were pelleted at 400 g for 6 minutes and resuspended in Staining Buffer (eBioscience, ThermoFisher) at a concentration of 3×10^6^ cells/mL. Aliquots of 200 µL (*i.e.*, 6×10^5^ cells) in 0.5 mL Eppendorf were used to prepare single stained controls and multi-colour samples for flow cytometry. Optimal volumes of antibodies were obtained from calculation of staining indexes, and each aliquot was stained with either single antibody (for single stained controls) or a cocktail of antibodies. For panels containing more than one Brilliant Violet (BV) dye, 30 µL of Brilliant Stain Buffer (ThermoFisher) was added to prevent dye aggregation. Cells were incubated at 4 °C in the darkness for 30 minutes, washed 2x with PBS (Gibco™, pH 7.4) and resuspended in 300 µL. Then, cells were stained with 1 µL of Fixable Viability Stain 660 (BD Bioscience, stock solution: 5 ng/µL in DMSO) at room temperature for 15 minutes, washed 2x in PBS and resuspended to a final volume of 400 µL in the flow tubes. For each experiment, we built a different panel of fluorescent markers to gate on cells subpopulations, which was compatible with our probe for PM order, namely Pro12A. *<u>Panel for PBMCs from healthy donors and patients</u>*: PBMCs were stained with anti-Human CD45-FITC (clone HI30, BioLegend), CD3-BV605 (clone SK7, BD Biosciences), CD8-PE-FIRE700 (clone SK1, BioLegend), CD14-BV711 (clone M5E2, BioLegend), CD19-PE-Cy7 (clone SJ25C1, BioLegend), CD56-APC-R700 (clone NCAM16.2, BD Biosciences), CD11c-PE (clone Bu15, BioLegend), FVS660 for cell viability. *<u>Panel for NK</u> <u>sorting and killing assay:</u>* Enriched NK cells were stained with CD3-PerCP-Cy5.5 (clone SK7, BioLegend), CD14-APC (clone M5E2, BioLegend), CD56-APC-R700 and FVS570 for cell viability. After incubation with K562 tumour cells, sorted NK cells were stained with CD56-PE (clone HCD56, BioLegend), CD3-PerCP-Cy5.5, CD107a-Alexa Fluor647 (clone H4A3, BD Biosciences) and FVS780 for cell viability. *<u>Panel for apoptosis study:</u>*PBMCs from healthy donors were stained with CD3-PerCP-Cy5.5, CD56-PE, CD19-PE-Cy7, Annexin-V-Alexa Fluor647 and FVS780 for cell viability. *<u>Panel for assessing the effect of freezing-thawing on</u> <u>PM order:</u>* PBMCs from healthy donors were stained with CD3-PerCP-Cy5.5, CD56-PE, CD19-PE-Cy7, CD11c-APC-R700 (clone 3.9, BD Biosciences), CD14-Alexa Fluor647 (clone M5E2, BioLegend), CD8-BV605 (clone SK1, BD Biosciences), Lactadherin-FITC (Prolitix) and FVS780 for cell viability. **Note:** Pro12A is generally measured using the BV421 and BV510 channels. Therefore, depending on the flow cytometer configuration, some commercial fluorochromes might not be suitable for coupling with Pro12A given their spectral spillover. In our flow cytometry setup (Fig. S19), the following dyes are not compatible with PM order measurement via Pro12A: BUV395, BUV496, BV421, BV480, BV510, AlexaFluor405. The fluorochromes BUV563 and BV570 might require additional optimization and spillover assessment given the spectral proximity with Pro12A detection channels. On the other hand, the following fluorochromes can be safely coupled with Pro12A for PM order measurement: BUV615, BUV661, BUV737, BUV805, BV605, BV650, BV711, BV786. In addition, Pro12A can be coupled with all fluorochromes from the PE and PerCP family as well as the other AlexaFluor derivatives and the red-excitable dyes. The panel of antibody-conjugated fluorochromes chosen for this study yielded negligible spillover into the channels occupied by the Pro12A signal used to measure membrane order, thereby ensuring robustness of the measurement (Fig. S20). Flow cytometry data were first gated on singlets, non-debris, and live (viability dye–negative) cells (Fig. S4). Subsequent gating identified each population as follows: Monocytes-derived Dendritic cells dim for CD11c (Mo-DCs (dim)): CD3⁻CD14⁺CD11c⁺; Monocytes-derived Dendritic cells bright for CD11c (Mo-DCs (bright)): CD3⁻CD14⁺⁺CD11c⁺; Dendritic cells dim for CD11c (DCs (dim)): CD3⁻CD14⁻CD56⁻CD19⁻CD11c⁺; Dendritic cells bright for CD11c (DCs (bright)): CD3⁻CD14⁻CD56⁻CD19⁻CD11c⁺⁺; B cells: CD3⁻CD14⁻CD56⁻CD19⁺; NK cells dim for CD56 (NK cells (dim)): CD3⁻CD14⁻CD19⁻CD56⁺CD8⁻; NK cells bright for CD56 (NK cells (bright)): CD3⁻CD14⁻CD19⁻CD56⁺⁺CD8⁻; CD8⁺ NK cells: CD3⁻CD14⁻CD19⁻CD56⁺CD8⁺; CD8⁻ T cells: CD3⁺CD14⁻CD56⁻CD8⁻; Cytotoxic T cells (CD8⁺ T cells): CD3⁺CD14⁻CD56⁻CD8⁺; NK-like T cells: CD3⁺CD14⁻CD56⁺CD8⁻; and CD8⁺ NK-like T cells: CD3⁺CD14⁻CD56⁺CD8⁺.

### Analysis of PM order

After antibodies incubation, cells (400 µL at a concentration = 1.5×10^6^ cells/mL) were stained with 1.5 µL of Pro12A (100 µM in DMSO) and vortexed vigorously. After 1 minute from staining, we acquired flow data for ∼5 minutes using a either a BD LSRFortessa 16-colour or a FACSCanto II flow cytometer. Before each experiment, we calibrated our flow pipeline by running DOPC-coated beads stained with Pro12A and adjusting the PMT voltages of the blue and green channels so to ensure the same median GP value for DOPC beads. Data from flow were analysed in FCS Express (DeNovo Software) and GP values from individual cells were normalized to the GP_DOPC_ to remove day-to-day variability. Specifically, before starting analysis on cells, we always ran freshly made BSLBs coated with DOPC membrane and measured their median GP value. This value was systematically subtracted from the GP value of individual cells to yield a normalised PM order value.

### NK cells sorting based on PM order

Before sorting, NK cells were enriched from PBMCs. Specifically, primary human NK cells were isolated by negative magnetic selection according to manufacturer protocol using a MojoSort human NK cell isolation kit (480054; BioLegend). The purity of enriched cells was >90%. Cells were counted (BioRad TC20) and checked for viability using trypan blue staining. After pre-enrichment, NK cells were stained with the antibodies cocktail (see above) and sorted based on their PM order. Cell sorting was performed on a BD FACSAria™ Fusion Flow Cytometer (BD Biosciences) equipped with 5 lasers, utilizing an 85 μm nozzle and running on FACS DIVA software. PM order was assessed by calculating the ratio between the intensities measured in the DAPI (ex. 405 nm, em. 450/50 nm) and the AmCyan (ex. 405 nm, em. 525/50 nm) channels. The DAPI/AmCyan ratio parameter was defined and created in the FACS DIVA software, and it was used as a regular fluorescence parameter thereafter. NK cell with high and low PM order were gated from the distribution of the DAPI/AMCyan ratios–as those falling below the 15th-percentile or above the 85th-percentile of the distribution–and collected in two falcon tubes. A small fraction of sorted cells was re-run both at the beginning and the end of the sorting procedure for purity check. Prior to sample analysis and sorting the instrument was calibrated with the BD® CS&T standard beads according to manufacturer’s instructions for optimal operation. Throughout the sorting, the sample tube’s temperature was kept at 20 °C to avoid changes in the cells’ membrane order. Sorted cells were kept on ice until plated in culture medium for functional assays or processed for RNA extraction.

### Migration assays on sorted NK cells

Migration assays were performed on agarose support. The protocol was adapted from a previously described method^106^. Briefly, 8-well polymer µ-slides (ibidi, 80821) were coated with recombinant human ICAM-1-Human-Fc chimera (RnD, 790-IC) at 3 µg/mL in PBS overnight at 4 °C. Slides were washed 3x with PBS, blocked with 2% bovine serum albumin (BSA) (Sigma-Aldrich) in PBS, and re-washed 3x. Agarose (0.5%, UltraPure Agarose, ThermoFisher) in RPMI+10% FCS-equivalent medium was prepared at 50 °C, and human CXCL12 (250 ng/mL RnD, 350-NS-010/CF) was added. 300 µL of agarose mixture was immediately pipetted in each µ-slide well. The agarose was placed at 4 °C for 1 h, and then transferred to 37 °C and 5% CO_2_ for 30 min. ΔGP-sorted NK cells were rested overnight at 37 °C and 5% CO_2_. The next day, cells were stained with CellTrace Violet (CTV, 1 µM, ThermoFisher) according to the manufacturer’s instructions. Cells were resuspended in >10 µL of RPMI+10% FCS containing propidium iodide (1.5 µM) and injected under agarose with a pipette. Cells were acclimatized at the microscope for at least 30 min at 37 °C and subsequently imaged every 15 s for 10 min using a Zeiss LSM780. Images were acquired with a Zeiss 20x (0.75 NA) objective, at 512 × 512 resolution in 8-bit using Zeiss Zen. Cells were tracked using the FIJI plugin TrackMate^107^. Propidium iodide-positive dead cells and tracks shorter than half the video length were excluded from analysis. One donor of the ten healthy donors collected yielded poor sample quality and was excluded from the migration assay.

### Killing assays on sorted NK cells

The killing capacity of NK-H and NK-L cells was assessed via degranulation assay as previously described^108^. Briefly, sorted NK cells were resuspended in complete PRMI medium (1,2×10^6^ live cells/mL), transferred in a V-bottom 96-well plates and co-cultured with K562 cells (ATCC) at 1:1 effector-to-target ratio in the presence of mouse anti-human CD107a Alexa Fluor 647 (BD Bioscience, clone H4A3, 2 µL/well) for 6 hours at 37°C. To block intracellular protein transport, BD GolgiStop (BD Bioscience) was added at a final concentration of 1:1000 during the assay after 1 hour of incubation. Once the 6 hours of incubation were completed, cells were washed twice in PBS + 2% FBS and stained with a cocktail of antibodies as described above for 30 minutes at 4°C in the dark, followed by two washes in PBS and staining with fixable Viability Stain (final concentration 1:1000) for 15 minutes at room temperature. Stained cells were washed in PBS once, stained with Pro12A and analysed in flow cytometry.

### Proximity Network Analysis

#### Sample preparation

Samples were processed using the Pixelgen Proxiome Kit (PROXIMM001) according to the manufacturer’s instructions. Briefly, samples were washed and fixed in 1% PFA, blocked, and then stained overnight at 4 °C with a pool of 155 barcoded antibodies and 3 mouse isotype controls. After staining, cells were washed and stabilized using a secondary antibody. Antibody barcodes underwent localized rolling circle amplification (RCA) of gap-filled padlock probes, followed by addition of linker oligonucleotides designed to hybridize to two proximal RCA products. Gap-fill ligation was then performed to incorporate the unique molecular identifier (UMI) and probe identifier (PID) sequences of the RCA products onto the hybridized linker oligonucleotide. One thousand cells per sample were transferred into a PCR amplification reaction to amplify the generated amplicons, followed by a second PCR to attach sequencing adapters. PCR products were purified using SPRIselect beads and quantified using the Qubit HS DNA assay. Purified products were diluted to 0.65 nM for Illumina P2 XLEAP kits or 0.49 nM for Illumina P3 and P4 XLEAP kits, spiked with 15% PhiX, and paired-end sequenced on an Illumina NextSeq 2000 using 44 cycles for Read 1 and 78 cycles for Read 2.

#### Data processing

PNA sequencing datasets were analysed using Pixelator, an open-source bioinformatics pipeline implemented in Nextflow via nf-core/pixelator (https://github.com/nf-core/pixelator) version 2.3.1. Briefly, raw sequencing reads were subjected to quality control and filtering before assembly into amplicons capturing protein interactions. Amplicons were subsequently grouped by marker assignment and error-corrected to generate an edge list representing the complete sample as a molecular graph. Leiden community detection was applied to resolve individual cells from this graph, retaining only components corresponding to intact cells for downstream analysis. Spatial metrics were then calculated for each cellular component and archived in PXL format along with associated metadata. Subsequent analyses were performed in R v4.4.3 using pixelatorR v0.16.0 (https://github.com/PixelgenTechnologies/pixelatorR) and Seurat v5.2.1. Data visualization was generated with ggplot2 v3.5.2 and ComplexHeatmap v2.22.0.

#### Analysis of sorted NK cells

Components were filtered based on multiple quality metrics: (1) molecule count thresholds of 25,000–100,000 UMIs per component; (2) normal Tau classification to exclude potential antibody aggregates^109^; and (3) isotype control antibody fractions (mIgG1, mIgG2a, mIgG2b) exceeding the sum of the global median and three times the median absolute deviation. Protein abundance data were normalized using centered log-ratio (CLR) transformation and scaled. Principal component analysis (PCA) was conducted, and the top 20 principal components were retained based on variance explained. Batch effects across samples were corrected by Harmony (v1.2.3) integration based on sample identity. Uniform Manifold Approximation and Projection (UMAP) was computed on Harmony-corrected dimensions for visualization. Cell clusters were identified by constructing a shared nearest neighbor (SNN) graph from Harmony-corrected dimensions, followed by Louvain clustering at resolution 0.1. Cluster-specific marker proteins were identified via Wilcoxon rank-sum tests. Additional cell annotation was performed by biaxial gating based on CD16 and CD56 CLR-normalized abundance levels. Differential protein abundance and proximity were assessed using pixelatorR functions RunDAA() and DifferentialProximityAnalysis(), respectively. For abundance testing, Wilcoxon rank-sum tests were performed on CLR-normalized values. Protein proximity was quantified using log2-ratios, representing the base-2 logarithm of the ratio between observed and randomly expected protein join counts^83^. For visualization, marker proteins were retained if detected in ≥0.2% of cells across ≥4 different samples within, with mean CLR abundance ≥1 and at least 3-fold higher than the combined isotype controls. Differential test results were displayed only for marker (pairs) with absolute effect size > 0.1. P-values were corrected for multiple comparisons using the Benjamini-Hochberg method, with significance thresholds at adjusted p < 0.05.

### UMAP analysis for clusters identification

Dimensionality reduction analysis was performed on live-CD45^+^-gated cells using FCS express software. All fluorescent parameters (except ΔGP) were scaled using a biexponential transformation setting the Below Zero parameter equal to 800. ΔGP values were scaled following a linear method. Cells were down sampled to ∼4×10^5^ cells (or 10^6^ for analysis on merged samples) following a random down-sampling method. For UMAP analysis, we set number of neighbours equal to 50, Min Low Distance equal to 0.4 and number of iterations equal to 600.

### RNA extraction and transcriptomic analysis

For the RNA extraction, a fraction of sorted NK cells was washed with 1X-PBS buffer 3 times, re-suspended in 1 mL TRIzol™ Reagent (15596018; ThermoFisher), and stored at −80°C until isolation. Organic extraction method was applied with chloroform addition and isopropanol precipitation. RNA Clean & Concentrator-5 (Zymo Research) was used for further purification and concentration. Three of the ten donors collected did not pass quality control, therefore we processed seven donors for transcriptomic analysis.

Library preparation and sequencing were performed at the Bioinformatics and Expression Analysis (BEA) core facility at Karolinska Institutet. All libraries were prepared using Takara’s SMART-SEQ V4 protocol and sequenced on a Nextseq 2000 sequencer with a P2 100 cycle flow cell. Cycling parameters were: 61 * 61 cycles paired end run with 8 cycles dual index. Base-calling and demultiplexing was performed using bcl2fastq (v2.20.0.422). Subsequently, quality control of the read sequences was performed with FastQC (v0.12.0, https://www.bioinformatics.babraham.ac.uk/projects/fastqc/), followed by preparation for alignment using CutAdapt (v3.5, adapter and quality trimming)^110^. Read alignment was completed with STAR aligner (v2.7.9a)^111^, made to the reference genome GRCh38 and genomic features were assigned using featureCounts (v1.5.1)^112^. Differential gene expression (DGE) analysis was performed with the DESeq2 package (v1.46.0)^113^ in R (v4.4.3)^114^, with the linear model design = ∼ Batch + GP_Status. The significance thresholds used were adjusted p value<0.05 (Benjamini-Hochberg adjustment), base mean>20 and absolute log2 fold change>0.3. For easy visualisation, principal component analysis (PCA) was performed using the pcaExplorer package (v3.0.0)^115^ in R (v4.4.3)^114^. Included heatmaps and plots were prepared using the pheatmap (v1.0.12, https://cran.r-project.org/package=pheatmap) and ggplot2 (v3.5.1) packages^116^ in R (v4.4.3)^114^.

Pathway analysis was performed as per a previously described protocol^117^. After the DGE analysis, RNK files were created and Gene-Set Enrichment Analysis (GSEA, v4.4.0)^118^ was performed for biological process (MSigDB v2026.1), molecular function (MSigDB v2026.1), and cellular component (MSigDB v2026.1), with a significance threshold cutoff of FDR<0.05. The scored gene ontology information for biological process was imported into Cytoscape (v3.10.4)^119^ and visualized using the EnrichmentMap (v3.5.0)^120^ app. The generated clusters with FDR adjusted p values (q values) were subsequently labelled using the AutoAnnotate (v1.5.2)^121^ app, to identify enriched pathways, that were differentially regulated. GSEA was also performed for KEGG pathways^122^ using the clusterProfiler package (v4.14.6)^123^ in R(v4.4.3)^114^. The enrichment of differentially expressed genes was also visualised in selected KEGG pathways^122^ using the pathview package (v1.46.0)^124^ in R (v4.4.3).

## Data availability

All data associated with this manuscript are publicly available in the FigShare repository (DOI: 10.17044/scilifelab.31882975) and in the Supporting Information.

## Code availability

Analysis of PM order from confocal microscopy images was performed our recently published open-source software VISION^104^. The analysis pipeline used to analyse transcriptomic data is available at the GitHub repository https://github.com/CSI-Nano-Lab/GP_RNAseq.

## Acknowledgments

This work is supported by Swedish Research Council Grants (grant no. 2020-02682, 2024-02993 and 2024-00289), Wellcome Leap’s Dynamic Resilience Program (jointly funded by Temasek Trust), Karolinska Institutet (2024-03250; 2024-03341; 2022-00803; 2020-00997), Cancer Research KI (2024-03488), SciLifeLab, Human Frontier Science Program (RGP0025/2022), Longevity Impetus Grant from Norn Group, Hevolution Foundation and Rosenkranz Foundation. We thank the SciLifeLab Advanced Light Microscopy facility and National Microscopy Infrastructure (VR-RFI 2016-00968) for their support on imaging. We also thank the core facility Bioinformatics and Expression Analysis (BEA) at NEO for their support with RNA extraction and analysis.

## Author contributions

L.A, E.S conceived and designed the experiments. L.A performed all the experiments together with Y.J and S.I (for experiment on PMBCs isolated from donors), P.S and V.C (for experiments on NK cells), L.B for experiments on NK cells migration, C.G and S.G for experiments on NK cells sorting, A.A for transcriptomic analysis, and F.R. for experiments on Proximity Network analysis. J.M and P.B. helped with collection of samples from long COVID patients and provide scientific insights. M.B and A.Ö helped with collection of samples from CLL patients and provided scientific insights. P.S provided scientific insights on NK cells killing assays. B.Ö provided scientific insights on NK cells. A.S.K. provided Pro12A and insights on its use. L.A and E.S co-wrote the manuscript. All authors read and corrected the manuscript.

