## Supplementary Figures for "Plasma membrane order maps functional diversity in immune cells"

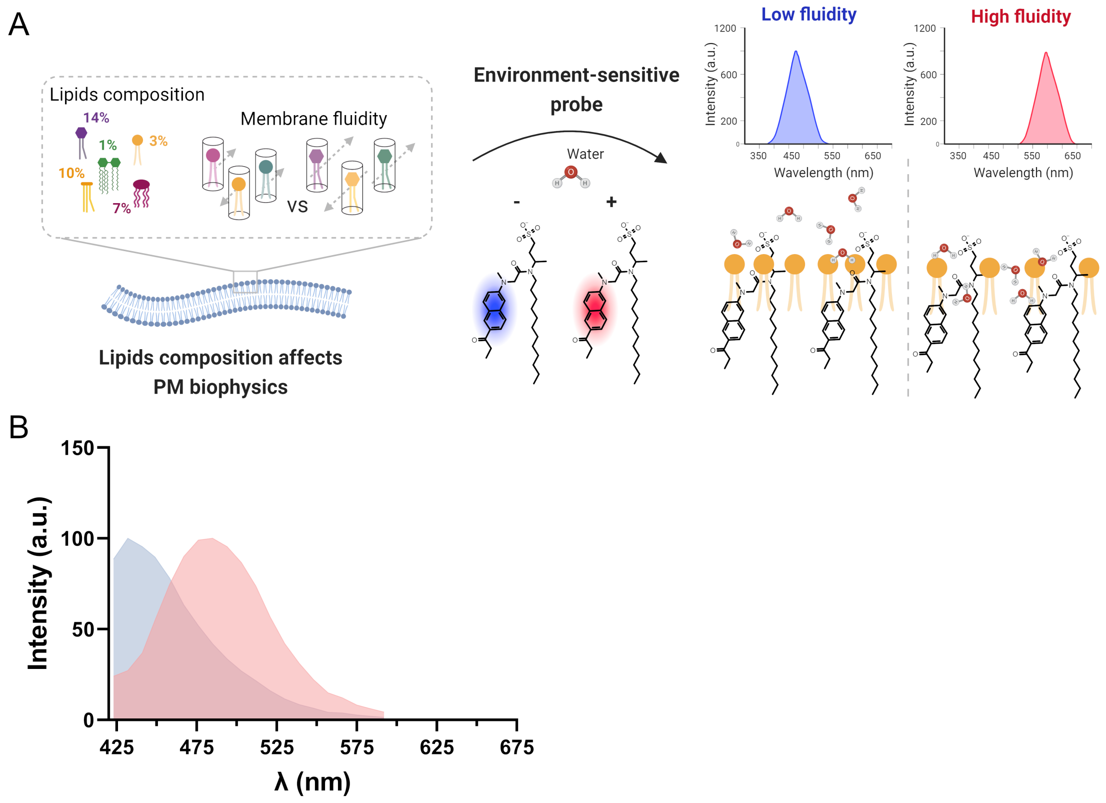


**Figure S1. A** Schematic of the mechanism for environment sensing for Pro12A dye. **B** Emission spectra of Pro12A staining silica beads coated with either low (blue curve, lipids composition = DPPC:Cholesterol 50:50%) or high (red curve, lipids composition DOPC 100%) order membranes.

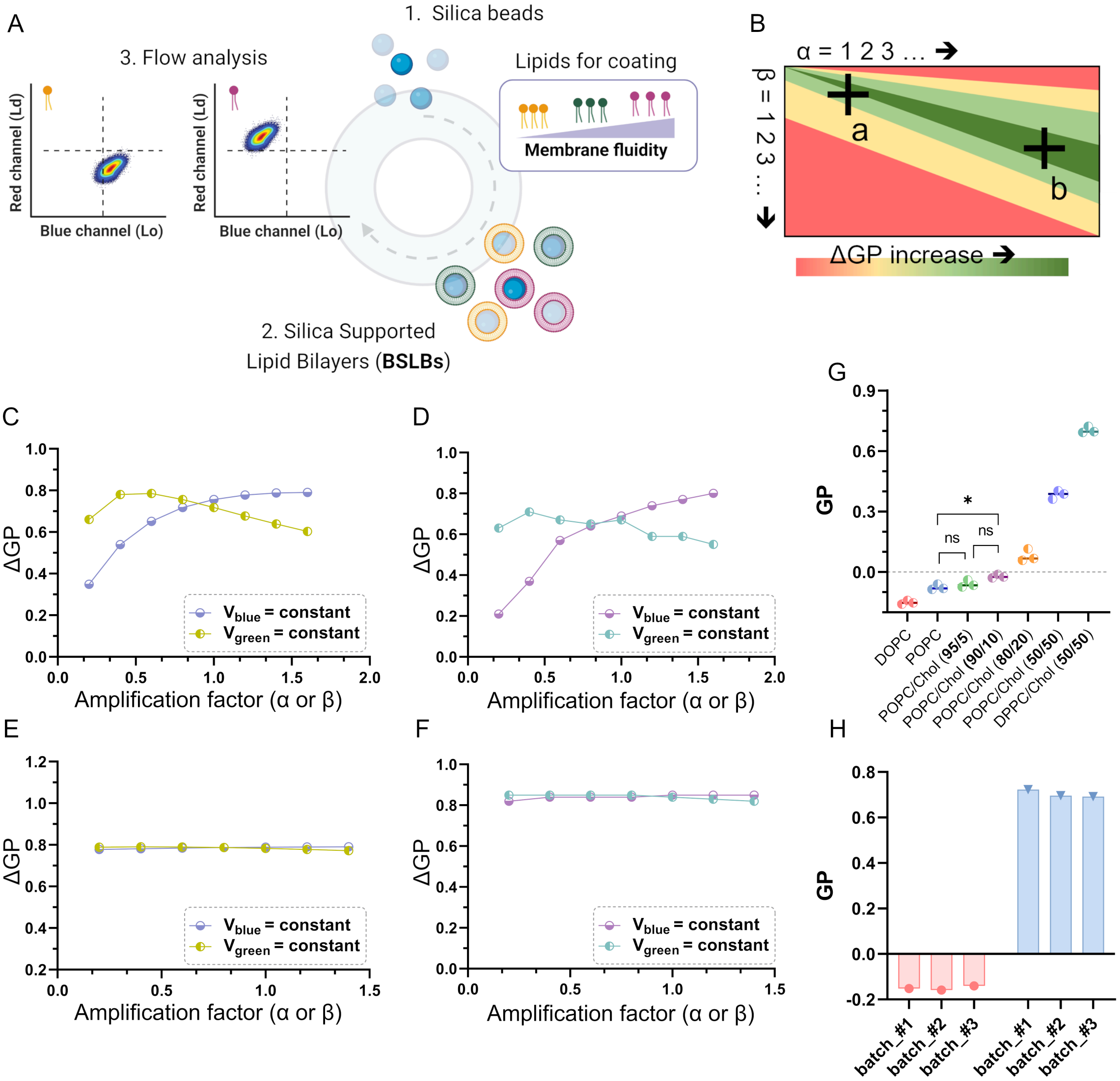


**Figure S2. A** Schematic of the synthesis of Silica Supported Lipid Bilayers (BSLBs) used for optimization of the flow setup. **B** Theoretical dependence of ΔGP on the amplification factors of the green (α) and blue (β) channels, calculated from the emission spectra of Pro12A in either low (DPPC:Cholesterol 1:1) or high (DOPC) order membranes. The black crosses indicate regions of constant high ΔGP at either low (a) or high (b) absolute intensities. **C-F** Theoretical (**C, E**) and experimental (**D, F**) ΔGP values as a function of the amplification factor within the region a (**C, D**) or b (**E, F**). **G** GP values from BSLBs with different membrane order (order increases from left to right). Each dot refers to the median GP value from one replicate. **H** GP values calculated from different batches of BSLBs coated with either low (DPPC:Cholesterol 1:1, blue bars) or high (DOPC, red bars) order membranes. Batches were prepared over ~270 days, demonstrating the reproducibility of BSLBs and usability as calibration beads.

The first step in our optimization process for PM order measurement was to assess both resolution and dynamic range of our flow cytometry setup by analysing model membranes with known membrane order. To do so, we prepared silica bead supported lipid bilayers (BSLBs) using pre-defined mixtures of lipids with different saturation degrees. BSLBs were stained with Pro12A and analysed in flow cytometry. Specifically, we collected the intensities in the blue (430-470 nm) and green (500-550 nm) regions of the emission spectrum and calculated the generalized polarization (GP) (*i.e.*, an experimental readout for membrane order) for individual beads. GP values were normalized to the most disordered membrane (i.e., DOPC-coated beads) and transformed into ΔGP values. Individual ΔGP values were defined according to the formula ΔGP = abs│$GPi_{-th}$ – $medianGP_{DOPC}$│(Eq. S1), where $GP_{i-th}$ and $medianGP_{DOPC}$ represent the membrane order of the *i-th* bead and the median GP of DOPC-coated beads, respectively (according to simplified version of Eq. S2).

When utilizing independent photomultiplier tubes (PMTs) detectors to measure intensities in the two channels, GP values can be potentially affected by different photons' amplification occurring at the two detectors. Therefore, we should consider two additional terms in the canonical equation for GP (se Eq. S1).

$$GP = \frac{(\alpha I_b - \beta I_g)}{(\alpha I_b + \beta I_g)} \quad \text{Eq. S2}$$

The *α* and *β* values represent the photons amplification factors in the blue and green channels, whereas $I_b$ and $I_g$ are the respective channel intensities. Using Eq. S2, we generated a theoretical 2D-map of ΔGP values (*i.e.*, $GP_{DPPCChol}$ – $GP_{DOPC}$, Fig. S2) as a function of pair-wise combination of *α/β* factors, starting from the emission spectrum of Pro12A in DOPC- and DPPC:Cholesterol 1:1-coated beads. Interestingly, the region of high ΔGPs (green band in Fig. S2B) widens with an increase in the amplification factors magnitude. Figures S2C-F show the theoretical and experimental ΔGP values obtained by systematically varying one amplification factor (*i.e.*, PMT voltage) at a time within the regions identified by the two black crosses in Fig. S2C. From our voltration experiments, we selected that pair of PMT voltages in the blue and green channels which yielded the highest and most stable ΔGP value.

As shown in Fig. S2G, using optimal PMT voltages ensured a wide dynamic range, enabling to detect differences in membrane with small changes in lipid order. Furthermore, given the stability and reproducibility of GP values from BSLBs over many days (Fig. S2H), we used 1,2-Dioleoyl-sn-glycero-3-phosphocholine (DOPC)-coated beads at the beginning of each experiment on cells, to calibrate our flow setup and normalize inter-day ΔGP measurements.

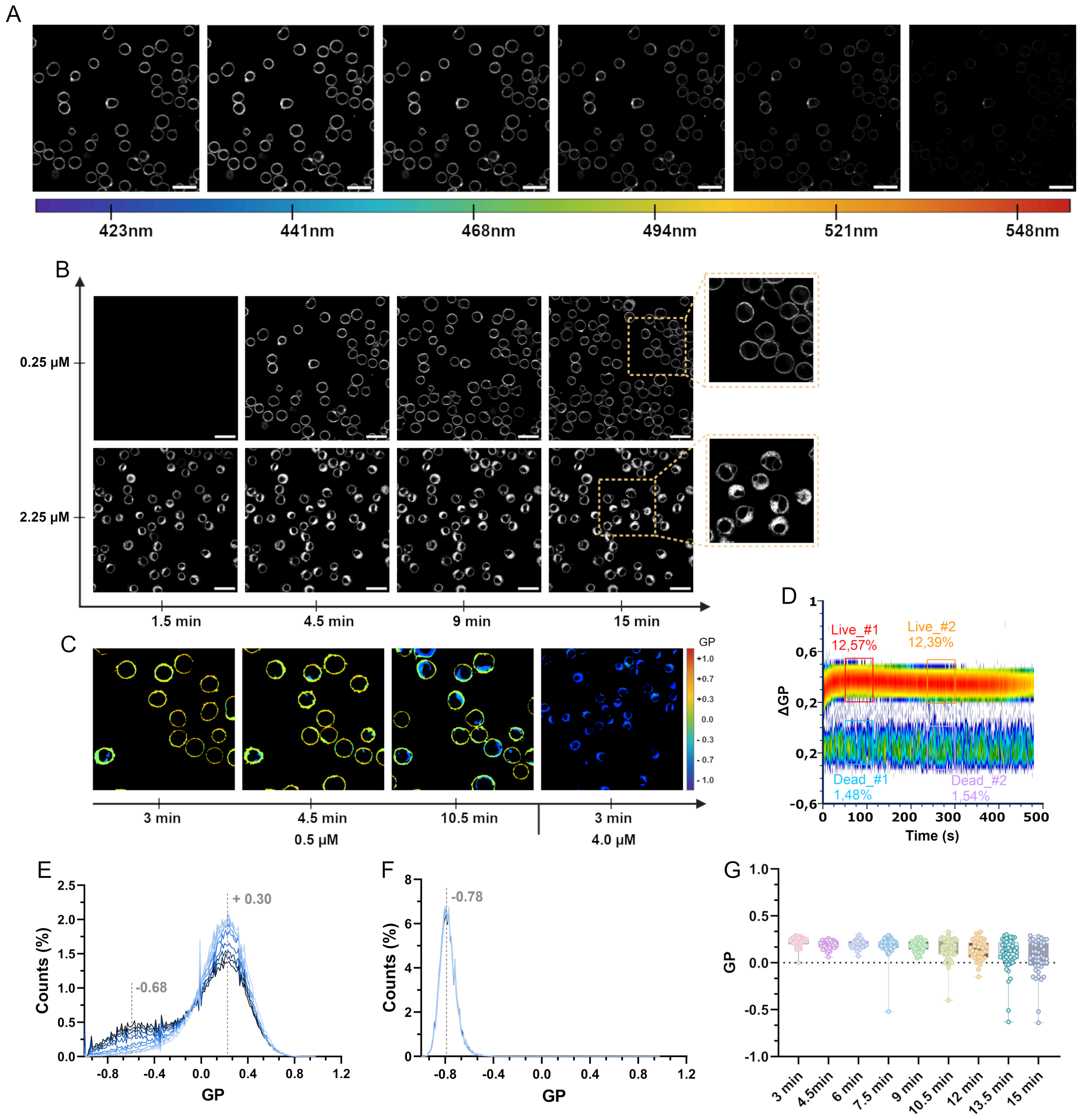


**Figure S3. A** Example of a λ-stack obtained with our spectral confocal microscope. Each image corresponds to a 9 nm-wide emission window. **B** Images showing the internalization of Pro12A in CEM cells as a function of time and at two different dye concentrations (i.e., 0.25 µM and 2.25 µM). **C** GP-colour coded images showing the shift in the emission of Pro12A (*i.e.*, GP value) between the plasma membrane and the intracellular environment, as a function of time and at two different dye concentrations. **D** Flow trajectory reporting ΔGP values over time in CEM cells**. E-F** Distributions of pixel-wise GP values as a function of time after staining (from light to dark blue curves) with either low (**E**) or high (**F**) concentration of Pro12A. The time lag between two consecutive curves is 3 minutes. The observed decrease in GP values (*i.e.*, increase of peak at -0.68 over time) is due to slow flipping of Pro12A to the inner leaflet of PM and, to a less extent, internalization of dye. At high probe concentration Pro12A becomes cytotoxic, and GP values decrease even further due to extensive internalization of dye and consequently staining of intracellular membranes in dead cells. **G** Single-cell GP median values as a function of time after staining, suggesting heterogeneity of PM permeability in CEM cells.

Next in the optimization protocol for the cytometry platform, we tested the compatibility of our pipeline with cell analysis. Pro12A is selective for the outer leaflet of PM but high concentrations might promote flipping to the inner leaflet and internalization[1]. To this end, we first screened different concentrations of Pro12A and staining times *via* spectral confocal microscopy on CEM cell lines (as model system for live cells), to identify optimal conditions for preventing internalization and cytotoxicity. By a proper titration of the dye, we were able to selectively stain the plasma membrane and prevent internalization up to ~20 minutes after staining (typical measurements of PM order in flow cytometry require ~5-6 min), thus demonstrating the greater performance of Pro12A compared to other commercially available probes (Fig. S3A-C)[1]. Lack of Pro12A internalization was also verified in our flow pipeline by analysing variations in ΔGP values over time (Fig. S3D). Indeed, following an initial stabilization time (due to the kinetic of Pro12A intercalation into the membranes), ΔGPs remained constant throughout the measurement. Furthermore, the negligible toxicity of our probe was confirmed by the unaltered ratio between the percentage of live and dead cells over time. Stability of ΔGP values over time was confirmed under confocal microscopy analysis by monitoring both distributions and cell-specific (CEM cell lines) medians of ΔGP values (Fig. S3E-G).

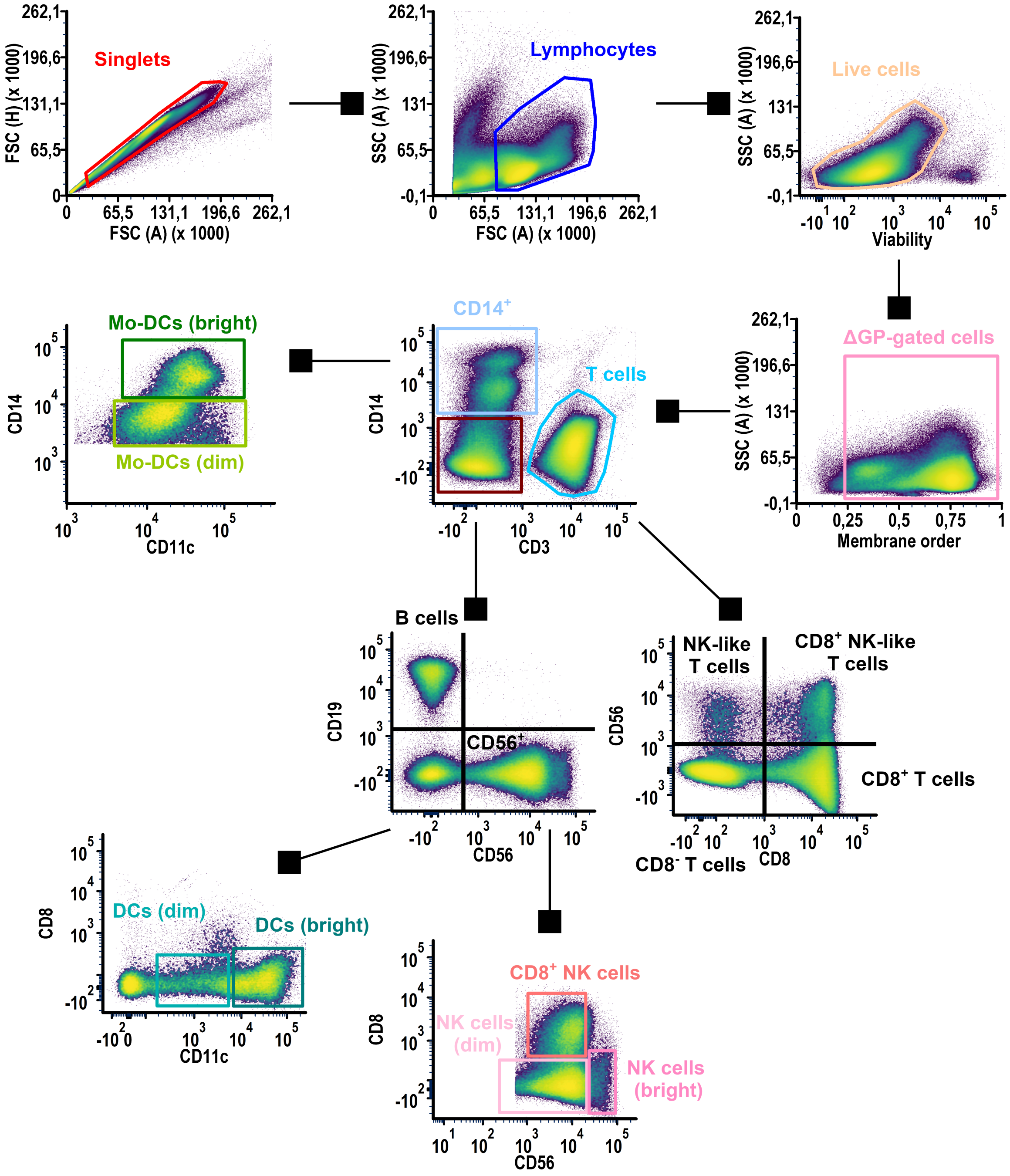


**Figure S4.** Gating strategy followed to identify the 12 subpopulations of immune cells from healthy and diseased patients. The example in the Figure S4 refers to pulled PBMCs isolated from healthy donors.

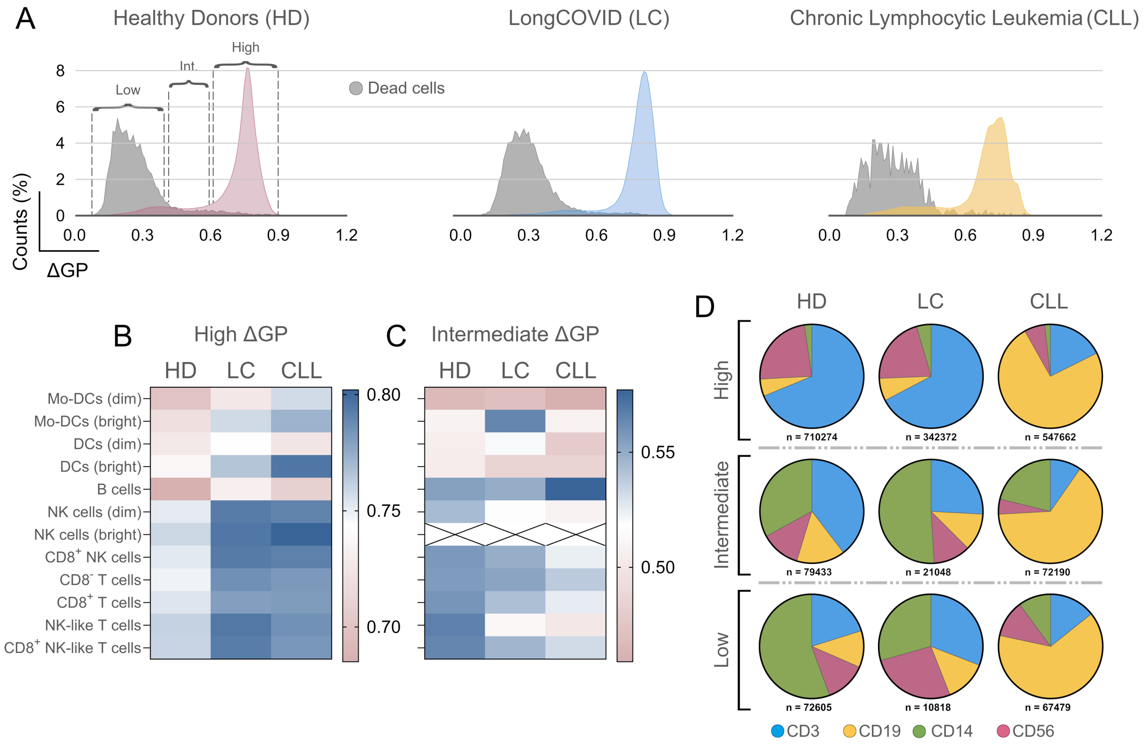


**Figure S5. A** PM order (ΔGP) distribution from pulled PBMCs from healthy and diseased donors. They grey distribution refers to the corresponding dead-gated cells. **B-C** Heatmaps showing variations in median ΔGP values for each immune cell subpopulation within the high ΔGP-gated and intermediate ΔGP-gated cells. **D** Pie charts showing the relative abundances of main immune cell types in each sample and within high, intermediate and low ΔGP-gated cells.

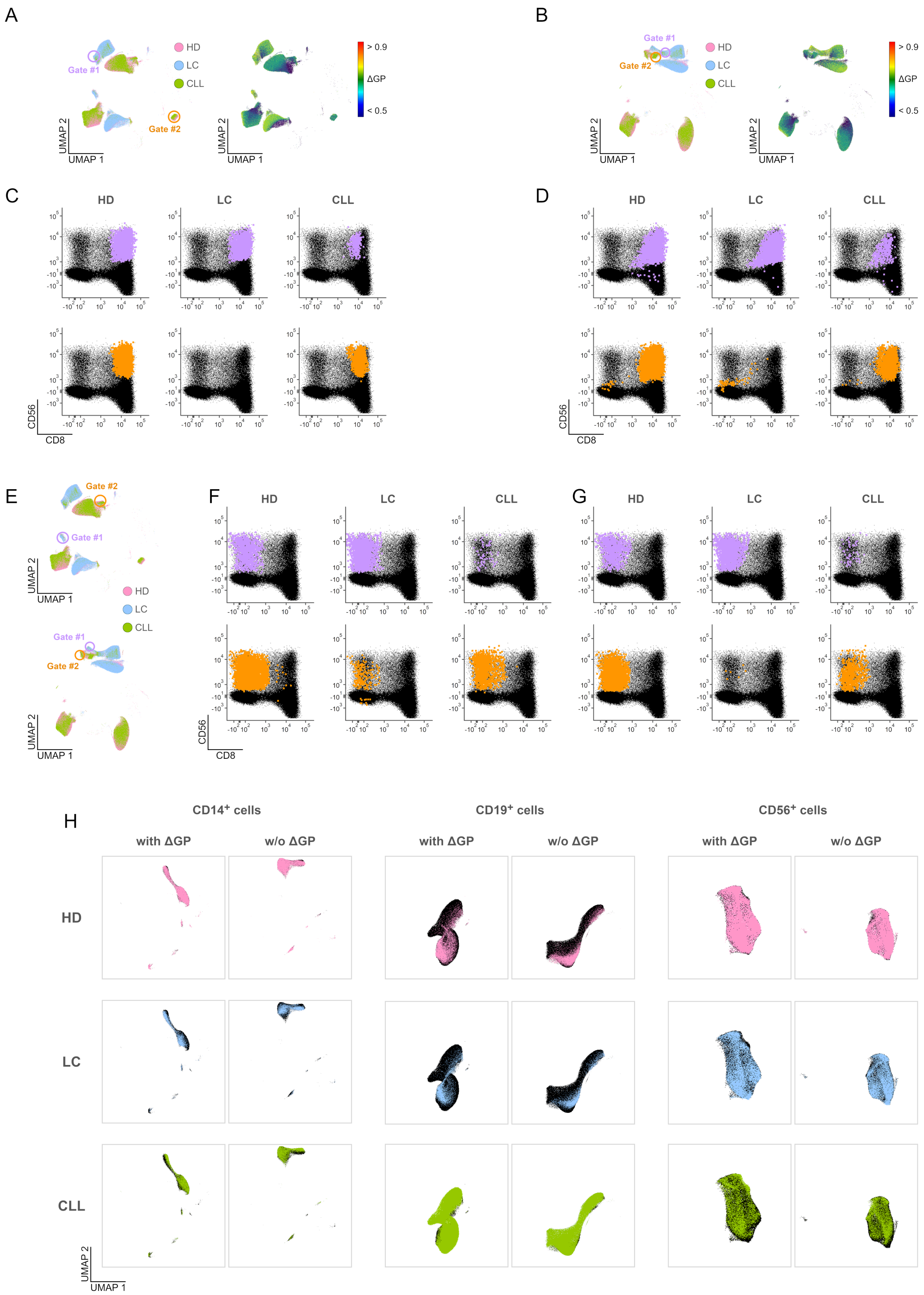


**Figure S6. A-B)** UMAP projections calculated from all samples–HD, LC and CLL–merged, selecting (**A**) or not (**B**) ΔGP as additional parameter during dimensionality reduction. The projections refer to T cell subpopulations and show the sample-specific clustering (left panel) and ΔGP value values (right panel). Gate #1 and Gate #2 in panel A and B refer to the same gated cells within T cells subpopulation (i.e., CD8⁺ NK-like T cells). **C-D)** 2D plots showing CD56 vs CD8 intensity distributions from cells in Gate #1 (purple dots) and Gate #2 (orange dots) for each patient sample, considering (**C**) or not (**D**) ΔGP as additional parameter in the UMAP analysis. **E)** UMAP projections calculated from all samples–HD, LC and CLL–merged, selecting (top) or not (bottom) ΔGP as additional parameter during dimensionality reduction. The projections refer to T cell subpopulations and show the sample-specific clustering. Gate #1 and Gate #2 in top and bottom panels refer to the same gated cells within T cells subpopulation ((i.e., NK-like T cells). **F-G)** 2D plots showing CD56 vs CD8 intensity distributions from cells in Gate #1 (purple dots) and Gate #2 (orange dots) for each patient sample, considering (**F**) or not (**G**) ΔGP as additional parameter in the UMAP analysis. H) UMAP projections for different immune cell types, considering or not ΔGP as additional parameter. Black dots refer to the total cells whereas each colour refer to a specific sample.

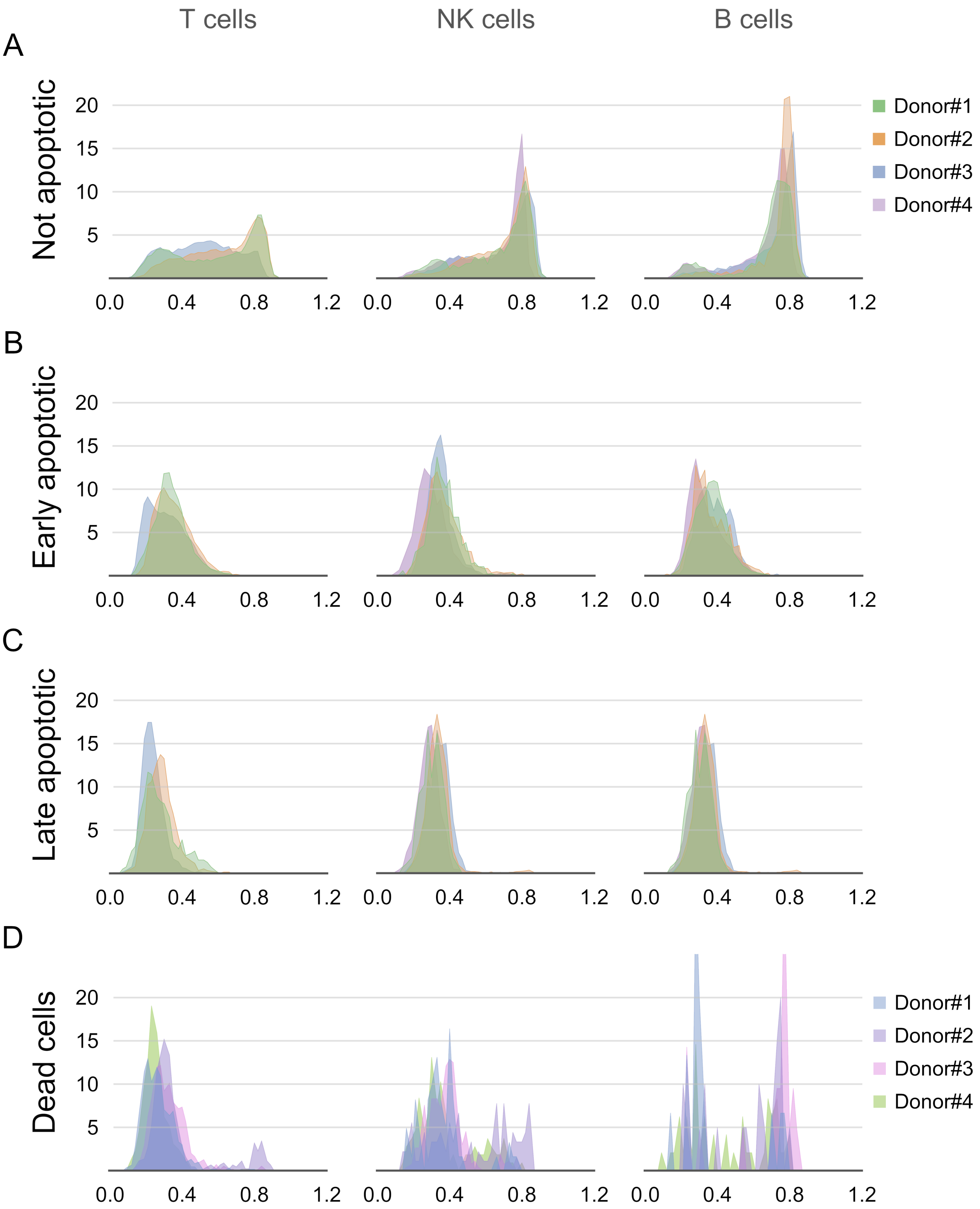


**Figure S7.** Apoptosis analysis on pulled PBMCs from four healthy donors. Plots show the overlap of PM order (ΔGP) distributions from individual donors for not apoptotic- (**A**), early apoptotic- (**B**), late apoptotic- (**C**) and dead-gated (**D**).

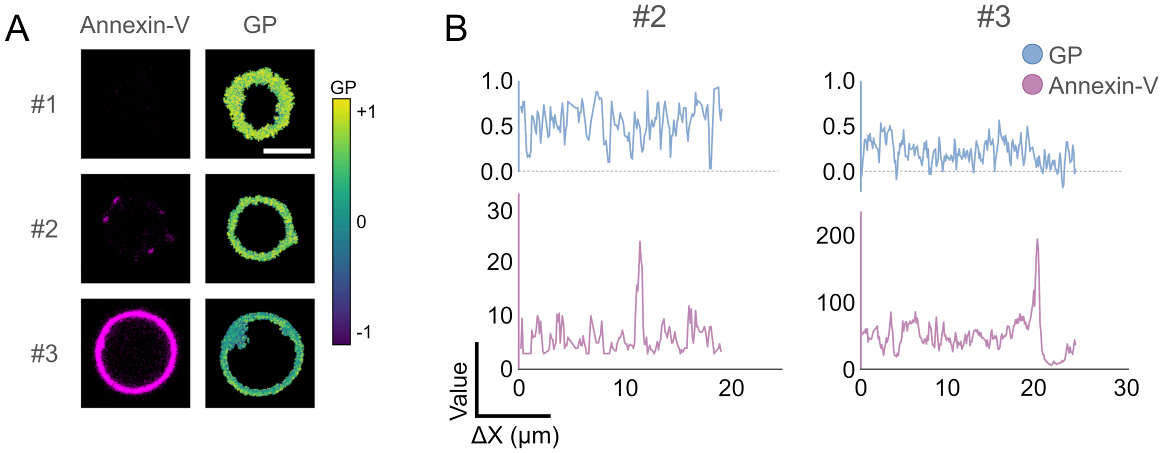


**Figure S8. A** Microscopy images acquired with a spectral confocal microscope, showing three different cells (#1, #2 and #3) after being stained with both Pro12A and Annexin-V-AF647. Magenta and pseudo-coloured images refer to Annexin-V and calculated GP values, respectively. The scale bar indicates 5 μm distance and the white arrows indicate Annexin-V puncta.

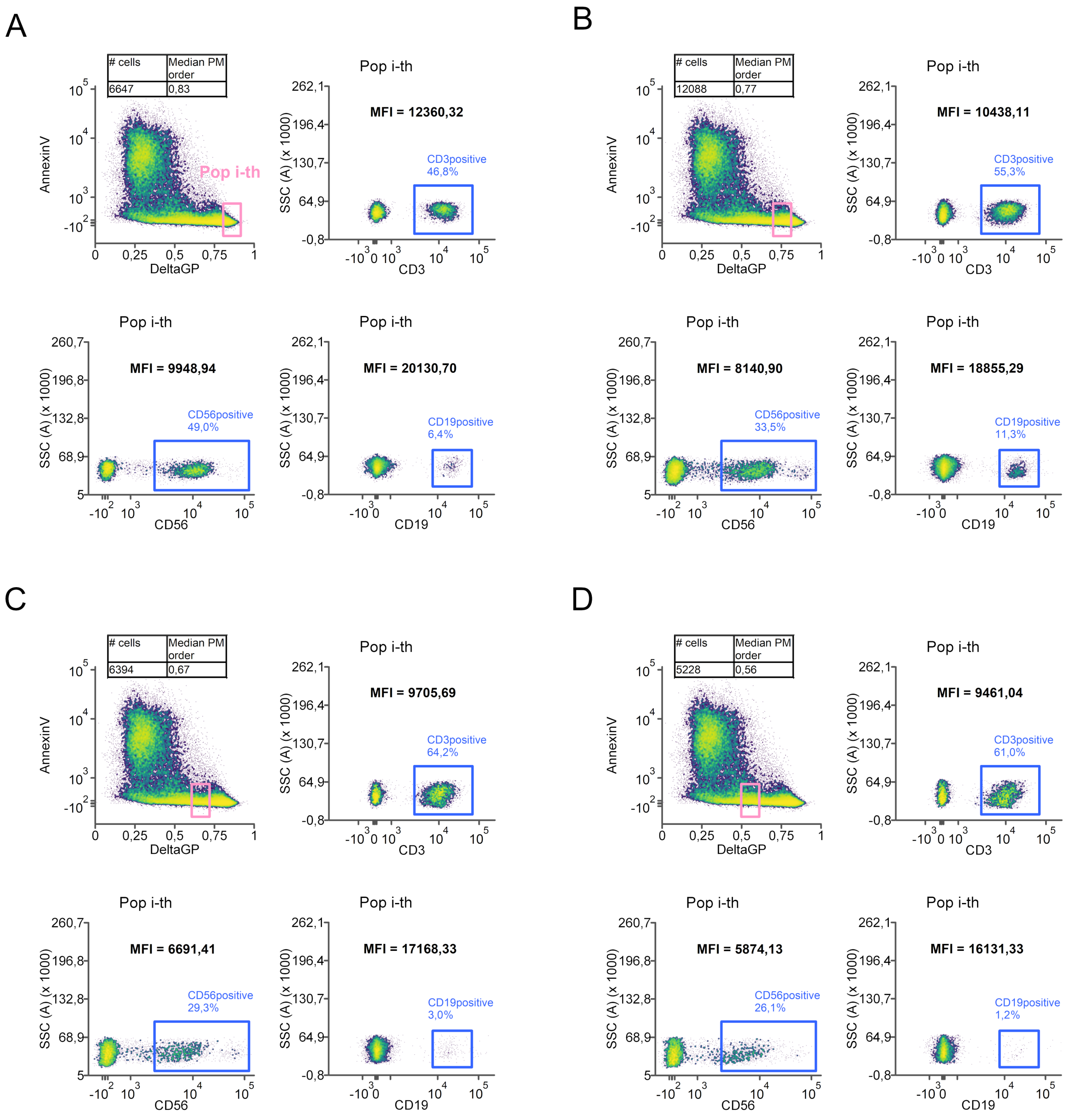


**Figure S9.** Expression level of main surface biomarkers for cell lineage in apoptosis studies. The expression of CD3, CD56 and CD19 was quantified from a subset of cells having median ΔGP equal to 0.83 (**A**), 0.77 (**B**), 0.67 (**C**) and 0.56 (**D**).

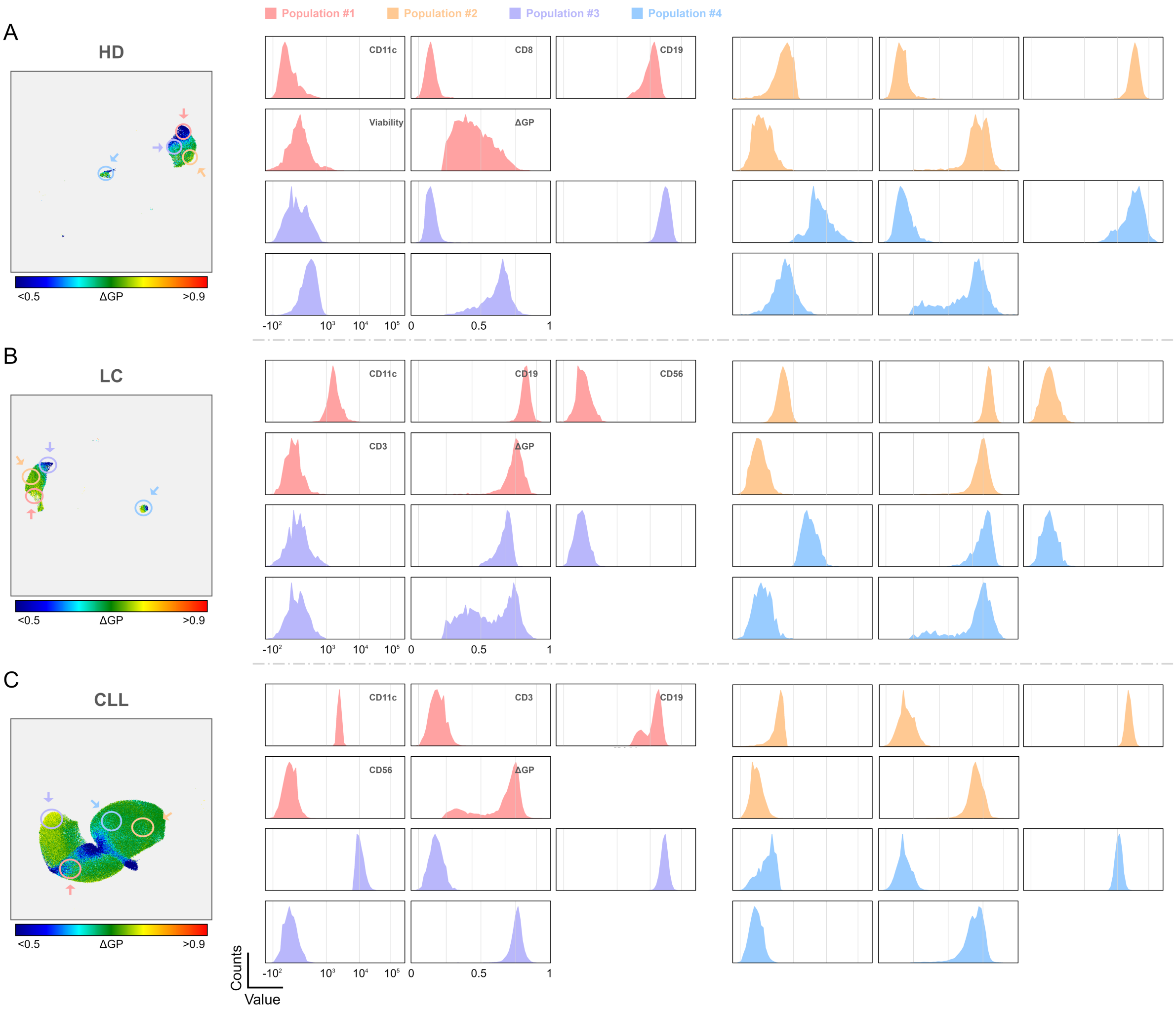


**Figure S10. A** UMAP projection (left box) and histograms showing the intensity distribution of different biomarkers and the distribution of ΔGP values (right panels). Plots refer to B cells ($CD19^+$) from healthy donors (**A**), Long COVID (**B**) and Chronic lymphocytic leukaemia (**C**) patients. Each coloured circle and corresponding arrow identify a different population within the B cells subset. The biomarker intensity histograms are rearranged in the same way across panels within the same sample.

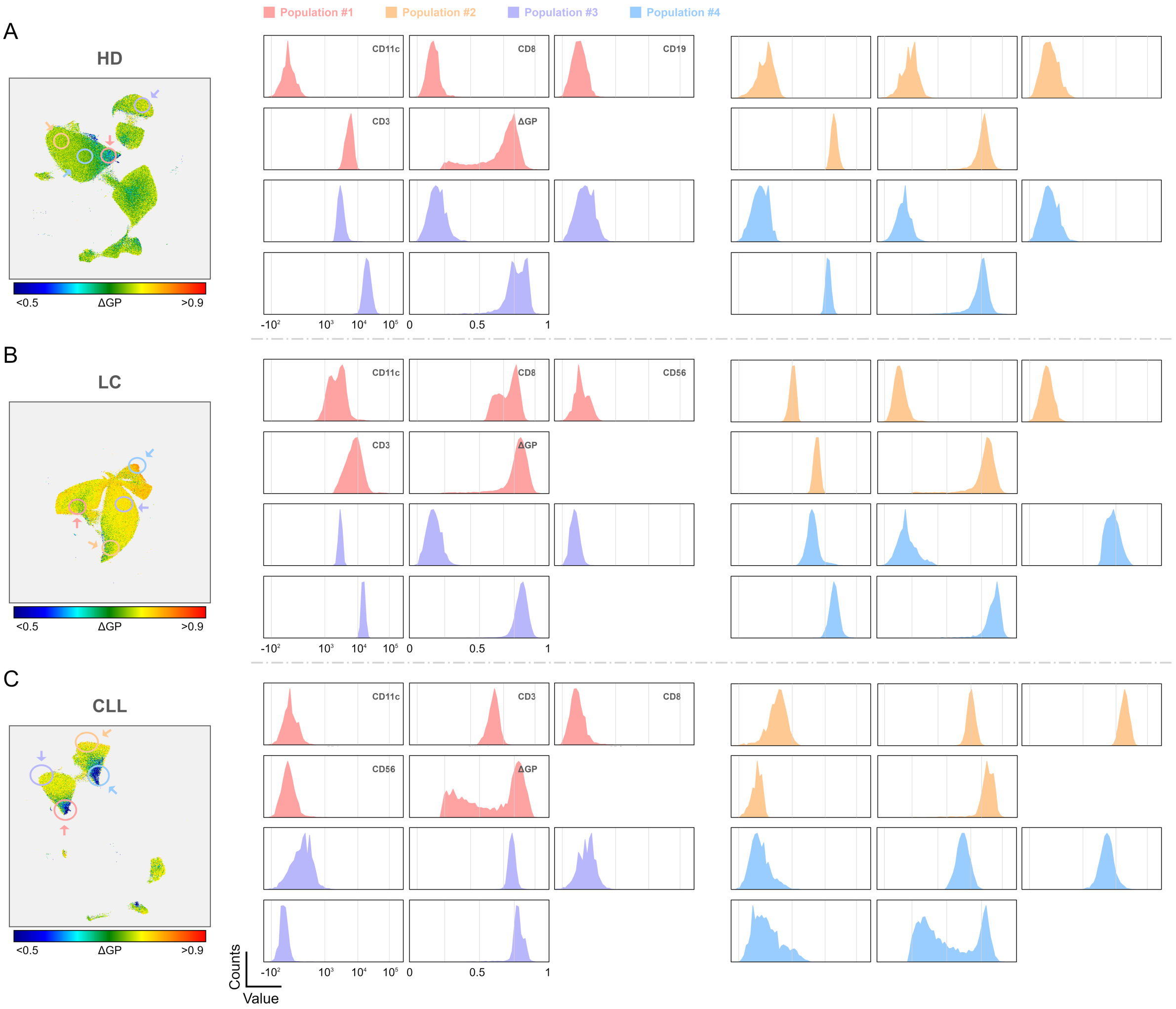


**Figure S11. A** UMAP projection (left box) and histograms showing the intensity distribution of different biomarkers and the distribution of ΔGP values (right panels). Plots refer to T cells ($CD3^+$) from healthy donors (**A**), Long COVID (**B**) and Chronic lymphocytic leukaemia (**C**) patients. Each coloured circle and corresponding arrow identify a different population within the T cells subset. The biomarker intensity histograms are rearranged in the same way across panels within the same sample.

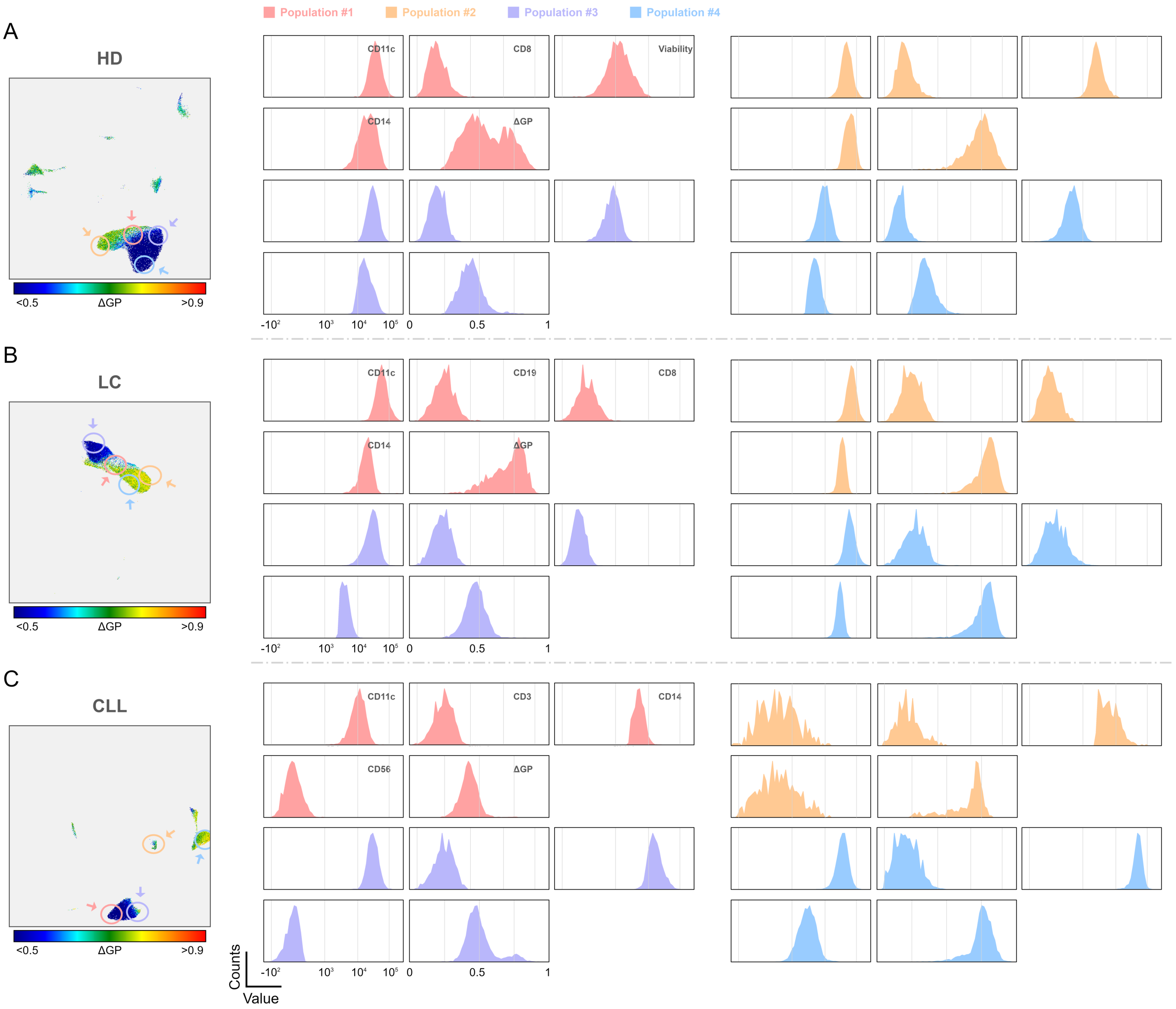


**Figure S12. A** UMAP projection (left box) and histograms showing the intensity distribution of different biomarkers and the distribution of ΔGP values (right panels). Plots refer to Mo-DCs cells ($CD14^{+}CD11c^{+}$) from healthy donors (**A**), Long COVID (**B**) and Chronic lymphocytic leukaemia (**C**) patients. Each coloured circle and corresponding arrow identify a different population within the Mo-DCs cells subset. The biomarker intensity histograms are rearranged in the same way across panels within the same sample.

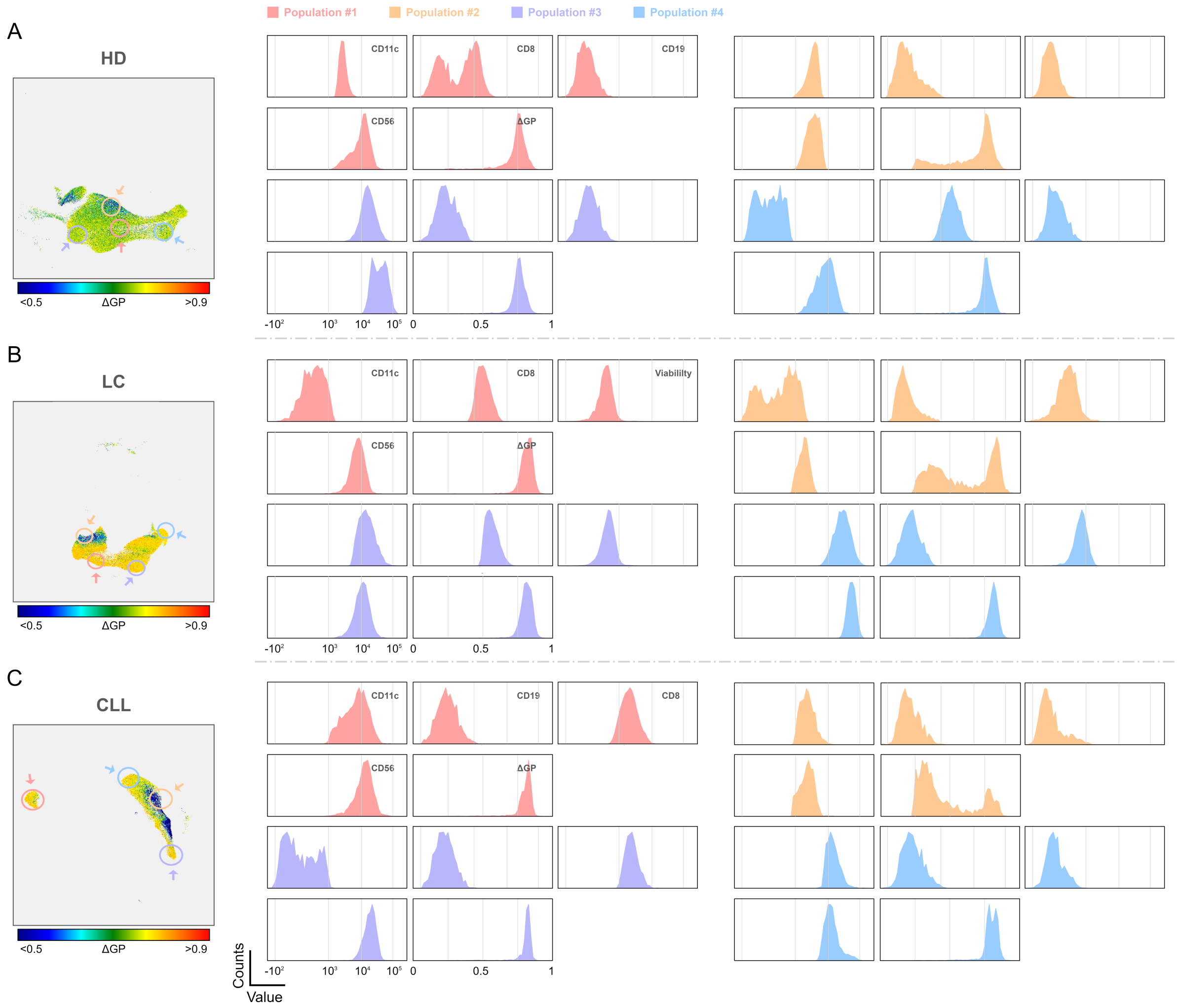


**Figure S13. A** UMAP projection (left box) and histograms showing the intensity distribution of different biomarkers and the distribution of ΔGP values (right panels). Plots refer to NK cells ($CD56^{+}$) from healthy donors (**A**), long COVID (**B**) and Chronic lymphocytic leukaemia (**C**) patients. Each coloured circle and corresponding arrow identify a different population within the NK cells subset. The biomarker intensity histograms are rearranged in the same way across panels within the same sample.

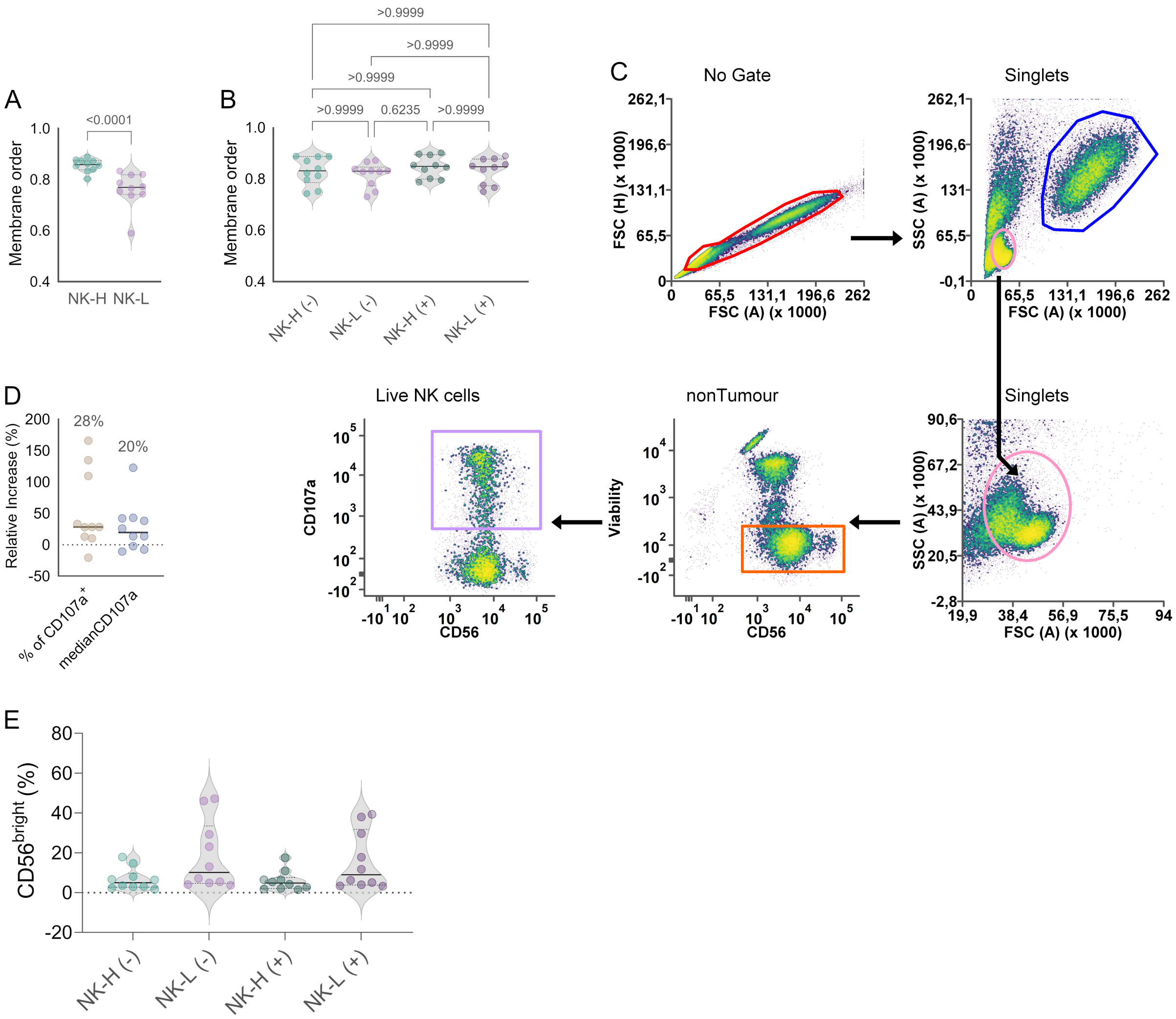


**Figure S14. A** Median membrane order values of NK cells from different healthy donors right after sorting. **B** Median membrane order values of sorted NK cells after 24 hours of recovery in cell medium, with (+) or w/o (-) incubation with K562 tumour cells. **C** Gating strategy to estimate number of live NK cells and degranulation percentage. **D** Relative percentage increase of CD107a expression level (intensity) and positive (CD107a$^{+}$) cells. **E** Violin plots showing the percentage of CD56$^{bright}$ cells within NK-H and NK-L subsets before (-) and after (+) exposure to tumour cells. In A and B, the small number represent the *p*-values calculated via the Kruskal-Wallis statistical test and the Dunn's test for multiple comparison. The threshold for statistical significance was set at $p$ = 0.05.

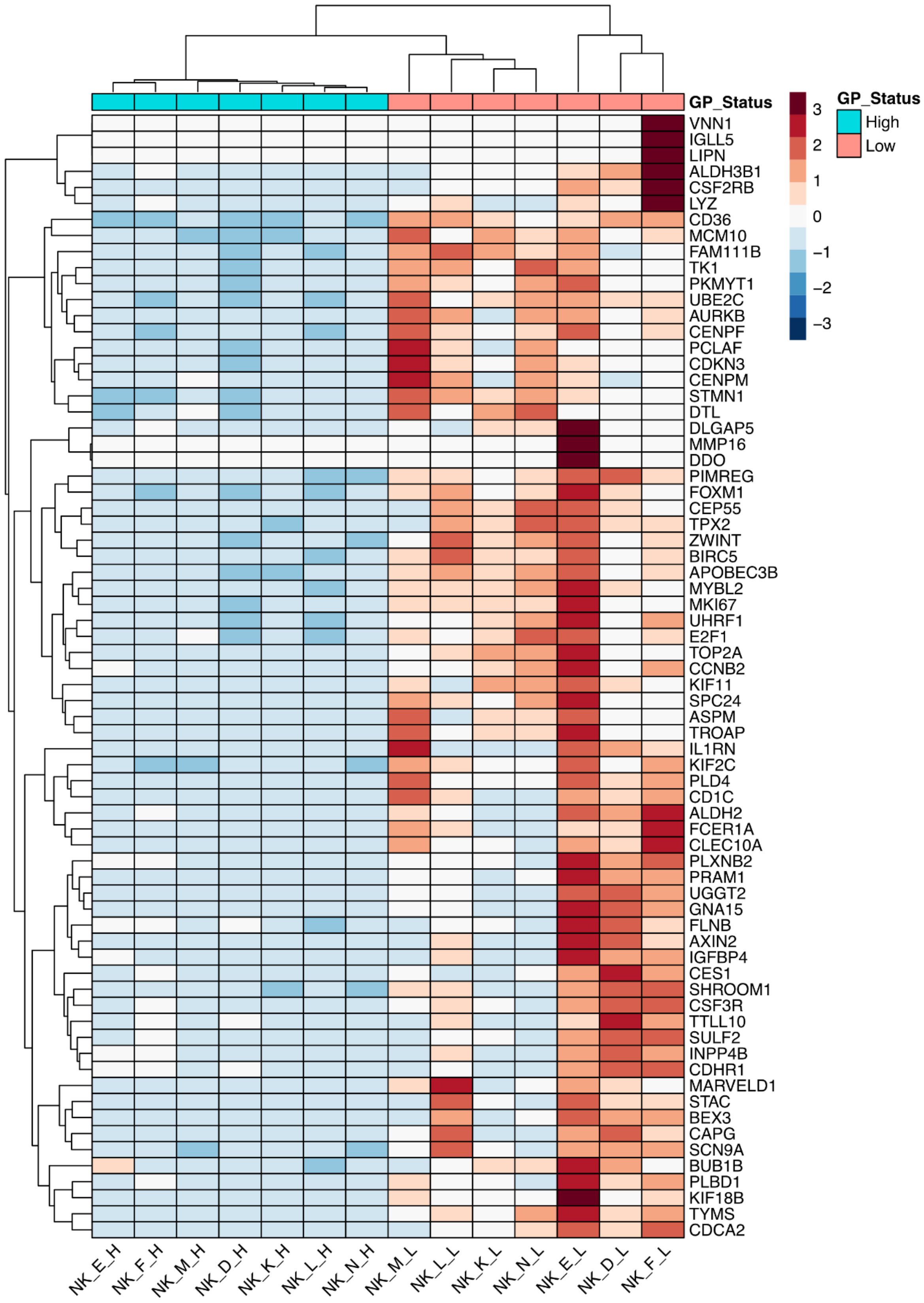


**Figure S15.** Heatmap showing the top 70 significant genes which were differentially expressed between NK-H and NK-L subsets.

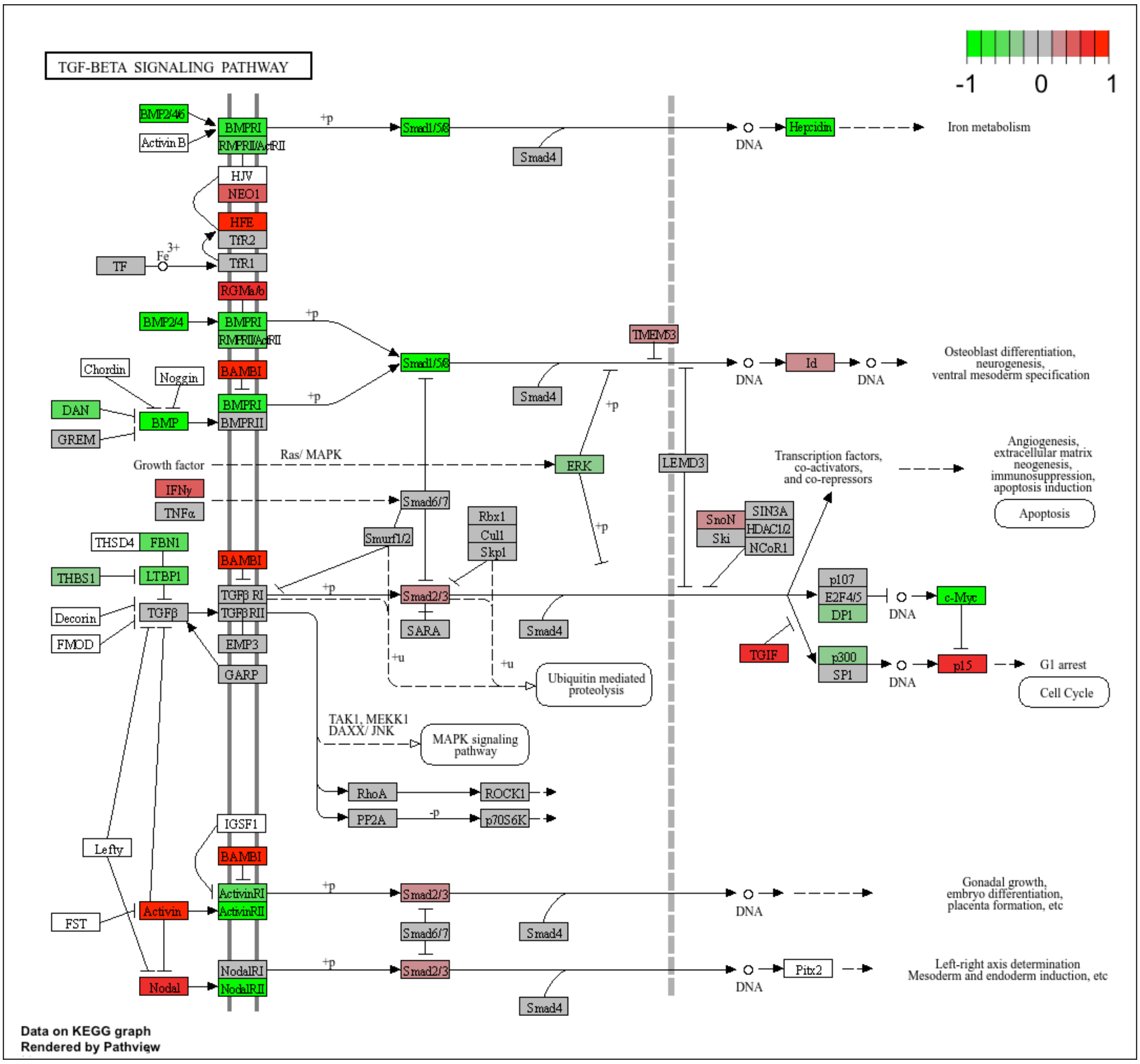


**Figure S16.** TGF-β signalling pathway from KEGG analysis, showing the location of identified genes.

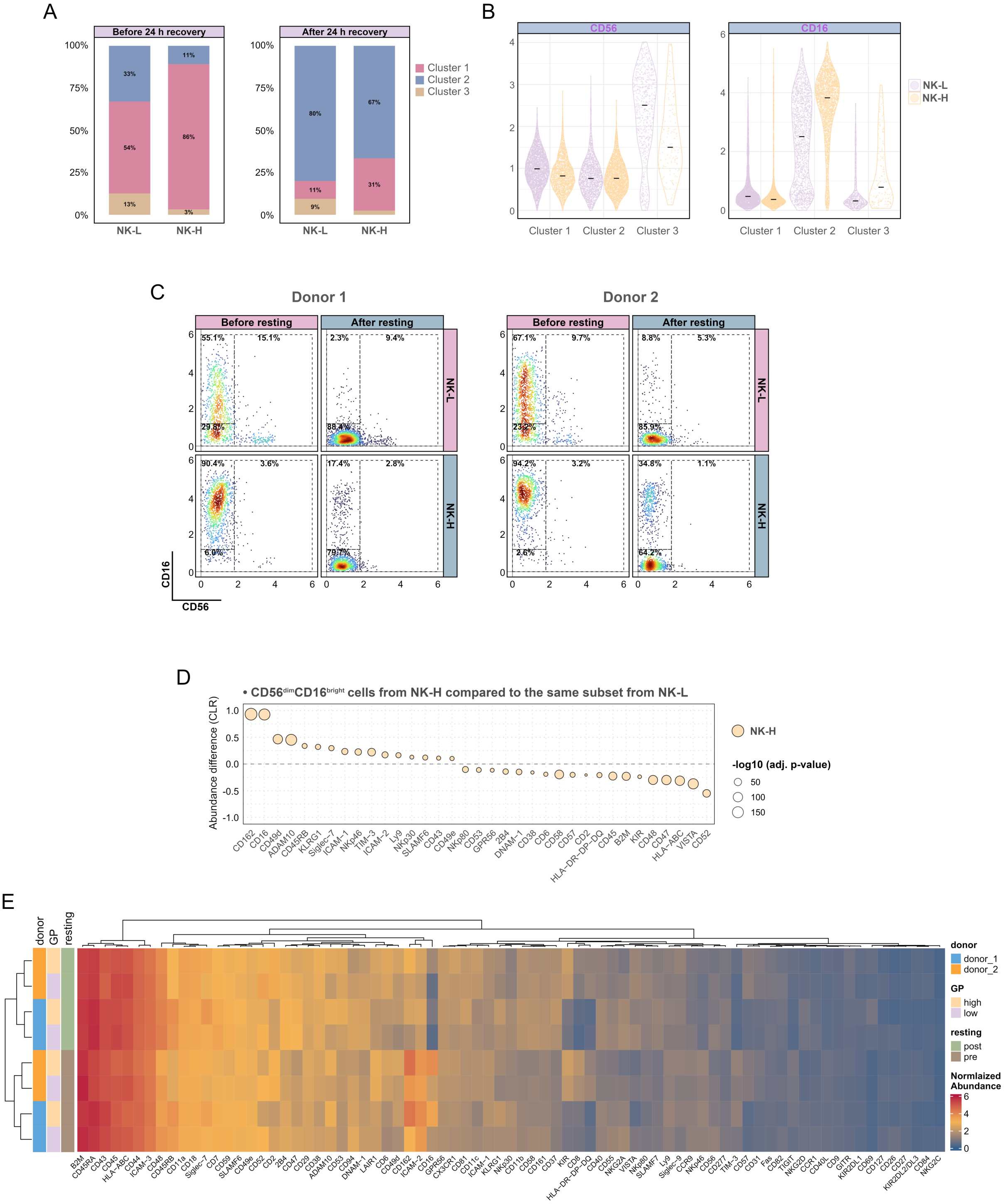

**Figure S17. A** Box plots showing the percentage composition of NK-H and NK-L in clusters 1, 2, and 3. **B** Violin plots showing the expression levels of CD56 and CD16 in NK-H and NK-L, and across clusters. **C** CD16 versus CD56 scatter plots as a function of resting time and PM order. **D** Differential abundance of surface markers between NK-H and NK-L subsets within the same phenotype (CD56$^{dim}$ CD16$^{bright}$). **E** Heatmap showing surface protein abundance from PNA analysis on sorted NK cells. Protein clustering is based on donor, PM order, and resting period conditions. In all panels, except panel C, data refer to pooled donors.

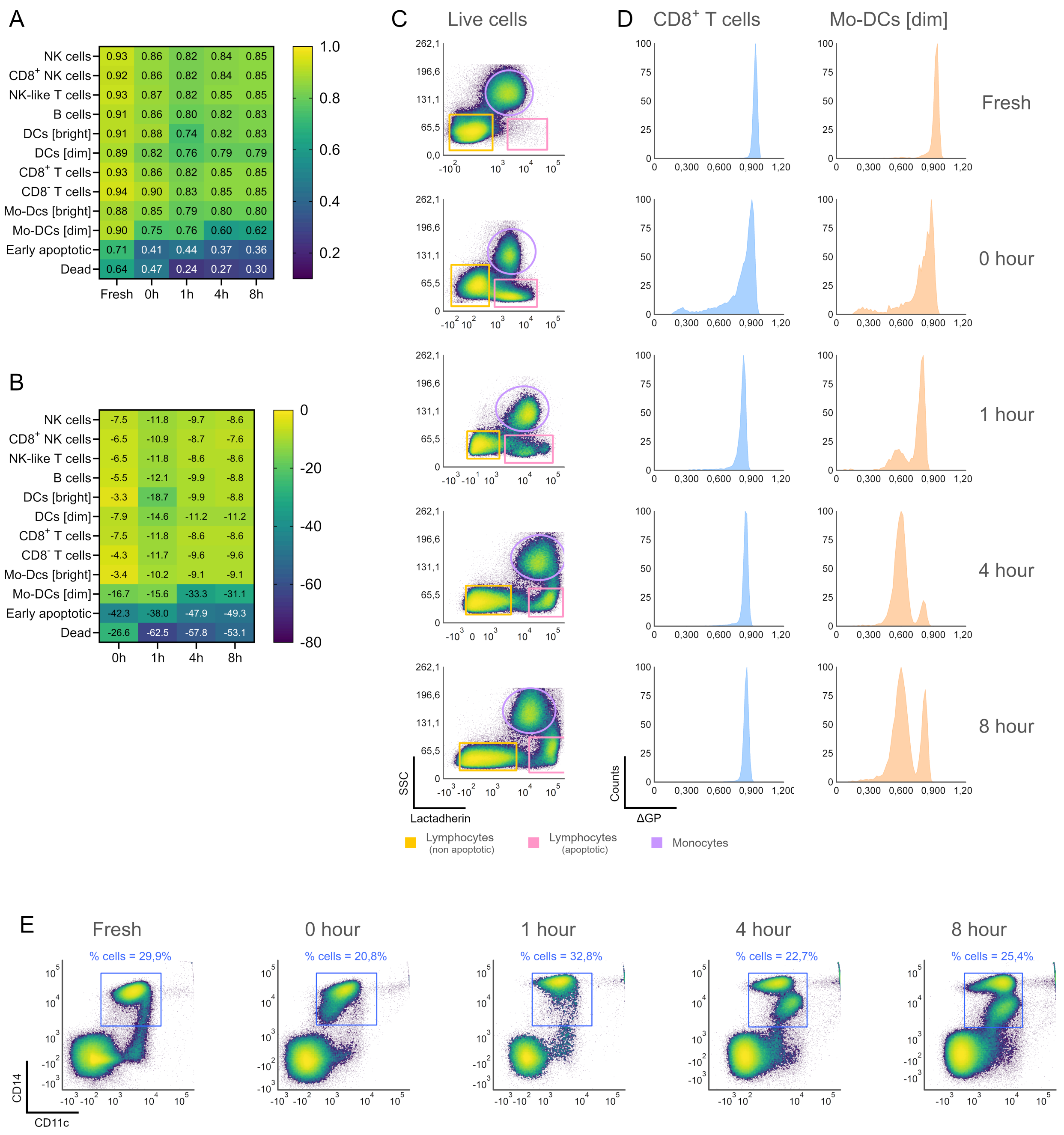


**Figure S18. A** Heatmap showing the median ΔGP values from PBMCs right after isolation (fresh) or after different post-thawing recovery times. **B** Relative decrease percentage of ΔGP values compared to freshly (no freezing) analysed PBMCs. **C-D** SSC versus Lactadherin plots (**C**) and ΔGP distributions (**D**) from freshly isolated PBMCs and cells after different post-thawing recovery times. E Percentage of $CD14^{+}$ cells from freshly isolated PBMCs or from frozen samples after different times in culture for post-thawing recovery.

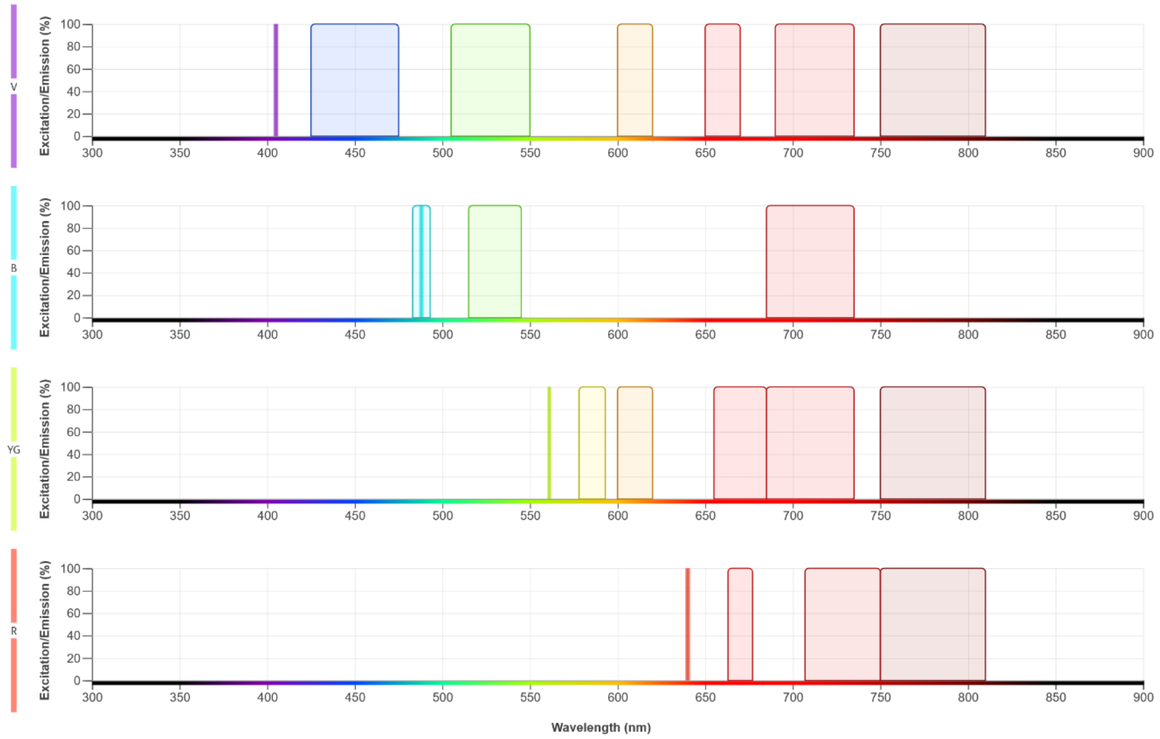


**Figure S19.** Configuration of our flow cytometer Fortessa BD LSRFortessa 16-colour, showing the lasers for excitation and the optical filters for wavelengths selection in the different channels.

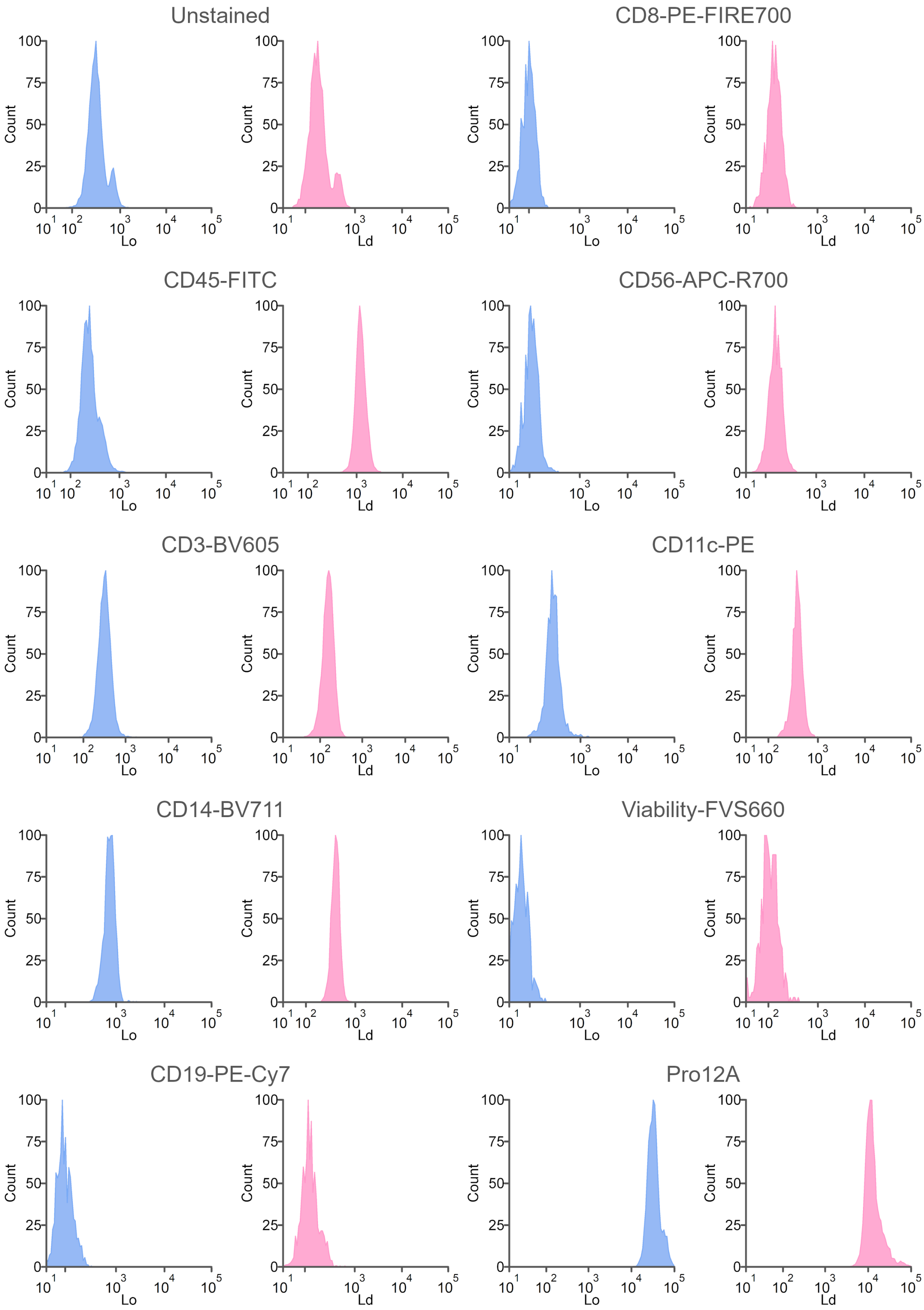


**Figure S20.** Spillover (i.e., intensity distribution) from antibody-conjugated fluorophores used to stain PBMCs from healthy donors and patients into the channels occupied by Pro12A signal. The *Lo* (ordered lipid phase) and *Ld* (disordered lipid phase) channels were used to measure membrane order.
